# PLD1-Dependent Regulation of Synaptic Integrity: Implications for Cognitive Resilience and Alzheimer’s Disease Pathogenesis

**DOI:** 10.64898/2025.12.08.692971

**Authors:** Sravan Gopalkrishnashetty Sreenivasmurthy, Shaneilahi M. Budhwani, Sanjana Mohanty, Marlene E Villarreal, Poonam Shah, Shyny K John, Chandramouli Natarajan, Karthik Ramaswamy, Kyle Sheth, Evan Wong, Phu Cuong Phan, Jose L Lopez, Satkarjeet Kaur Gill, Joshua Currie, Klarissa Garza, Agenor Limon, Thomas A Green, Balaji Krishnan

## Abstract

Dysregulation of Phospholipase D1 (PLD1) has been implicated in the progression of neurodegenerative diseases, including Alzheimer’s Disease (AD). This study investigated PLD1 signaling in synaptic integrity and cognition during aging and in a late-onset AD mouse model, hypothesizing differential effects of PLD1 modulation on synaptic vulnerability. AAV2-mediated gene transfer was employed to overexpress (PLD1 OXP) or attenuate (PLD1 ATT) PLD1 in aged wild-type (WT) and 3xTg-AD mice. To validate these constructs, differentiated PC12 cells were utilized. Within these cells, a model of post-mitotic neurons, PLD1 OXP exhibited a notable reduction in both the average neurite length and the percentage of neurite-bearing cells. This suggests that elevated PLD1 activity exerts a significant influence on neurite outgrowth. Conversely, PLD1 ATT did not inhibit neurite formation, indicating it is not detrimental at the cellular level. These cellular findings paralleled in vivo observations - electrophysiological studies revealed PLD1 OXP impaired long-term potentiation (LTP) and synaptic transmission, particularly in aged WT mice, whereas PLD1 ATT improved synaptic function in 3xTg-AD mice. Behaviorally, PLD1 ATT enhanced spatial working memory and reduced anxiety-like behavior, notably in 3xTg-AD mice. These results highlight that tight PLD1 regulation is vital for maintaining synaptic integrity and cognitive resilience. Thus, PLD1 attenuation may serve as an important complement to immunotherapeutic approaches by strengthening synaptic resilience in neurodysfunctional states, including AD and related dementia (ADRD).

## INTRODUCTION

Phospholipase D (PLD) is a conserved lipolytic enzyme superfamily found across diverse species, from bacteria to humans (Frohman 2015; Bowling et al. 2020). This enzyme catalyzes the hydrolysis of phosphatidylcholine (PC) to generate phosphatidic acid (PA) and choline, key signaling molecules in cellular processes (Brown et al. 2017). In higher vertebrates, multiple PLD isoforms exist, each possessing distinct protein-protein interaction domains and subcellular localizations, highlighting their diverse roles in signal transduction. These isoforms are found at the plasma membrane (PLD1, PLD2) (Zhang et al. 2004; Barber et al. 2022), endosomes, endoplasmic reticulum (ER, PLD1) (Nakagawa et al. 2017), Golgi (PLD1) (Riebeling et al. 2009), and lysosomes (PLD3) (Nackenoff et al. 2020). Notably, only PLD1 and PLD2 exhibit direct lipolytic activity on PC, while the specific functions of other isoforms are still being elucidated. PLD1 and PLD2, while generating PA from PC (McDermott et al. 2020), possess distinct structural features, regulatory mechanisms, and subcellular distributions that underpin their specialized roles. Both isoforms contain conserved catalytic HKD motifs within domains II and IV (McDermott et al. 2020). However, they differ in other domains, including Phox (PX) and Pleckstrin Homology (PH) domains, which mediate protein-protein interactions and lipid binding, influencing their regulation and membrane association (Park & Han 2018). Notably, human PLD1 contains a variable loop region, absent in PLD2, which appears to confer autoinhibition, as its deletion increases basal activity (McDermott et al. 2020). Consistent with this, PLD1 typically exhibits low basal activity and requires stimulation by upstream signals, including protein kinase C-α (PKCα), ADP ribosylation factor (ARF), and Rho family GTPases, for robust activation (Park & Han 2018). In contrast, PLD2 is often considered to be more constitutively active (Vitale et al. 2001) and is frequently localized to the plasma membrane (Yin et al. 2010). PLD1, however, exhibits a more dynamic distribution where it is predominantly found in intracellular compartments, including the Golgi complex, early and late endosomes, lysosomes, the endoplasmic reticulum (ER), and perinuclear vesicles (Park & Han 2018). Upon cellular stimulation, a pool of PLD1 translocate to the plasma membrane. The stimulus-dependent translocation of PLD1 underscores its role as a regulated signaling node, poised to integrate diverse upstream signals and generate localized PA pools precisely where and when needed for downstream effects, such as modulating membrane dynamics during exocytosis or endocytosis (Dall’Armi et al. 2010), and this includes regulation of synaptic neurotransmission (Humeau et al. 2001; Vitale et al. 2001).

While PLD1 plays constructive roles in neuronal development and function, compelling evidence, significantly driven by prior work from this laboratory, points towards a detrimental role for aberrantly elevated PLD1 levels in the context of aging and Alzheimer’s Disease (AD). We established clinical relevance by discovering aberrant overexpression of PLD1, but not PLD2 in crude synaptoneurosomal fractions from post-mortem AD patients hippocampal tissues compared to age-matched controls (Krishnan et al. 2018). This observation was corroborated by findings of upregulated PLD1 immunoreactivity in reactive astrocytes surrounding amyloid plaques in AD brains (Jin et al. 2007) and supported by historical reports of generally elevated PLD activity in AD (Kanfer et al. 1996; Higgins & Flicker 2000). Consistent with such human data, we identified an age-dependent increase in the crude synaptoneurosomal fractions of hippocampal PLD1 in the late-onset 3xTg-AD mouse model (Oddo et al. 2003; Javonillo et al. 2022), which recapitulates key aspects of both amyloid-β (Aβ) and tau pathology (Krishnan et al. 2018), where this widely used transgenic model harbors human transgenes for mutant Amyloid Precursor Protein (APP_Swe_), Presenilin-1 (PS1_M146V_), and microtubule-associated protein tau (MAPT_P301L_) (Barber et al. 2023). Taken together, these observations in human clinical samples and mouse models of AD/ADRD points to an important role for PLD1 in progressive neurodegeneration by impinging on synaptic integrity.

Small molecule-mediated attenuation of PLD1 prevented AβO- and TauO-induced synaptic and memory deficits in C57BL/6 mice (Krishnan et al. 2018). Extending these findings, chronic treatment paradigms using the specific and well-tolerated PLD1 small molecule inhibitor VU0155069 (VU01) demonstrated efficacy in promoting synaptic and cognitive resilience in 3xTg-AD mice. In mice treated during earlier stages of pathology (starting at 6 months), pharmacological repeated PLD1 attenuation prevented the expected decline in cognitive function (Novel Object Recognition, NOR and Fear Conditioning, FC) and maintained better synaptic function compared to vehicle-treated controls (Bourne et al. 2019). Similar protective effects were observed even when treatment was initiated at a later stage (starting at ∼11 months), when significant Aβ and tau pathology is established (Natarajan et al. 2023). In these older mice, repeated VU01 treatment (a) ameliorated deficits in NOR and FC, (b) preserved both high frequency stimulation long term potentiation (HFS-LTP) and low frequency stimulation long term depression (LFS-LTD) at Schaffer collateral (SC) hippocampal CA3-> CA1 synapse, and (c) maintained dendritic spine integrity in the CA1 region, characterized by preservation of mature mushroom spine features and filamentous spine length. These collective findings strongly suggest that aberrantly elevated PLD1 significantly increases the vulnerability of synapses to the pathological insults, characteristic of the aging and AD brain.

Despite the established roles of PLD1 in development (McDermott et al. 2020) and the clear benefits of inhibiting pathologically elevated PLD1 in AD/ADRD (Krishnan et al. 2018; Bourne et al. 2019; Natarajan et al. 2023), a critical knowledge gap remains regarding the consequences of modulating PLD1 levels after brain development is complete. Specifically, the impact of post-developmental elevation of PLD1 on synaptic function and vulnerability, particularly within the context of normal and pathological aging, has not been directly investigated. While developmental studies suggest potentially neuroprotective roles, the lack of an overt cognitive deficit in 6-month-old C57Bl/6 mice despite PLD1 attenuation by repeated VU01 treatment (Suppl. Fig. 1) and the detrimental association of aberrantly elevated PLD1 with AD pathology raises the question of how PLD1 upregulation or downregulation, independent of its developmental functions, contributes to age-related synaptic decline.

This leads to the central question motivating the present study: Is PLD1 signaling intrinsically involved in setting the threshold for synaptic vulnerability during the aging process? More specifically, does artificially increasing PLD1 levels after development exacerbate age-related synaptic dysfunction or vulnerability to neuropathological insults? Conversely, does attenuating PLD1 levels during aging promote resilience, even in the absence of overt AD pathology? Addressing these questions is crucial for validating the hypothesis that elevated PLD1 causally contributes to synaptic vulnerability in the aging brain.

To directly address the role of post-developmental PLD1 modulation in synaptic vulnerability, a strategy combining in vitro proof-of-concept studies with in vivo functional assessments using targeted gene manipulation in aging mouse models was employed. AAV vectors (serotype 2), a powerful and widely adopted tool for gene delivery and manipulation within the CNS (Chan et al. 2017; Kanaan et al. 2017) was engineered to drive either the overexpression of human PLD1 cDNA or the attenuation of endogenous PLD1 expression via a validated short hairpin RNA (shRNA) sequence (Zhu et al. 2012; Luo et al. 2017), alongside appropriate control vectors. PC12 cells were first differentiated using nerve growth factor (NGF) to induce neuronal-like morphology. Following differentiation, cells were transiently transfected with either PLD1 cDNA (for overexpression), PLD1-targeting shRNA (for knockdown), or appropriate control constructs. The efficacy of PLD1 overexpression or silencing were subsequently confirmed by assessing any observable effects on neurite morphology. The in vitro validation was corroborated with the in vivo validation using aging WT (C57Bl/6 or 129Sv) or 3xTg-AD mice where multimodal functional assessments were performed using behavior, electrophysiology, and biochemistry to understand PLD1 as a key determinant influencing synaptic integrity and functional outcomes during aging, thereby impacting the progression towards neurodysfunctional states including Alzheimer’s disease and related dementias (AD/ADRD).

## RESULTS

### PLD1 Modulation Alters Neurite Number and Length in Differentiated PC12 Cells

PLD1-modulating constructs were validated in vitro using PC12 cells (derived from a rat pheochromocytoma of neural crest origin (Tominami et al. 2024)). Upon treatment with Nerve Growth Factor (NGF), these cells cease dividing and differentiate into a sympathetic neuron-like phenotype, extending neurites and expressing neuronal markers (Taglialatela et al. 1991; Taglialatela et al. 1992). Differentiated PC12 cells thus offer a simplified, yet relevant, post-mitotic neuronal context to perform proof-of-concept experiments.

PC12 cells were treated with NGF (100 ng/mL) for 7 days with low-serum media to induce differentiation that was observed as neurite outgrowth and branching (Fig. 1A & Suppl. Fig. S2 D, E, F). These differentiated cells were then transiently transfected for 48 hours with plasmid constructs encoding either Control (eGFP/SCR), ATT (PLD1 shRNA), or OXP (PLD1 OXP) (Fig. 1B).

**Figure 1:**
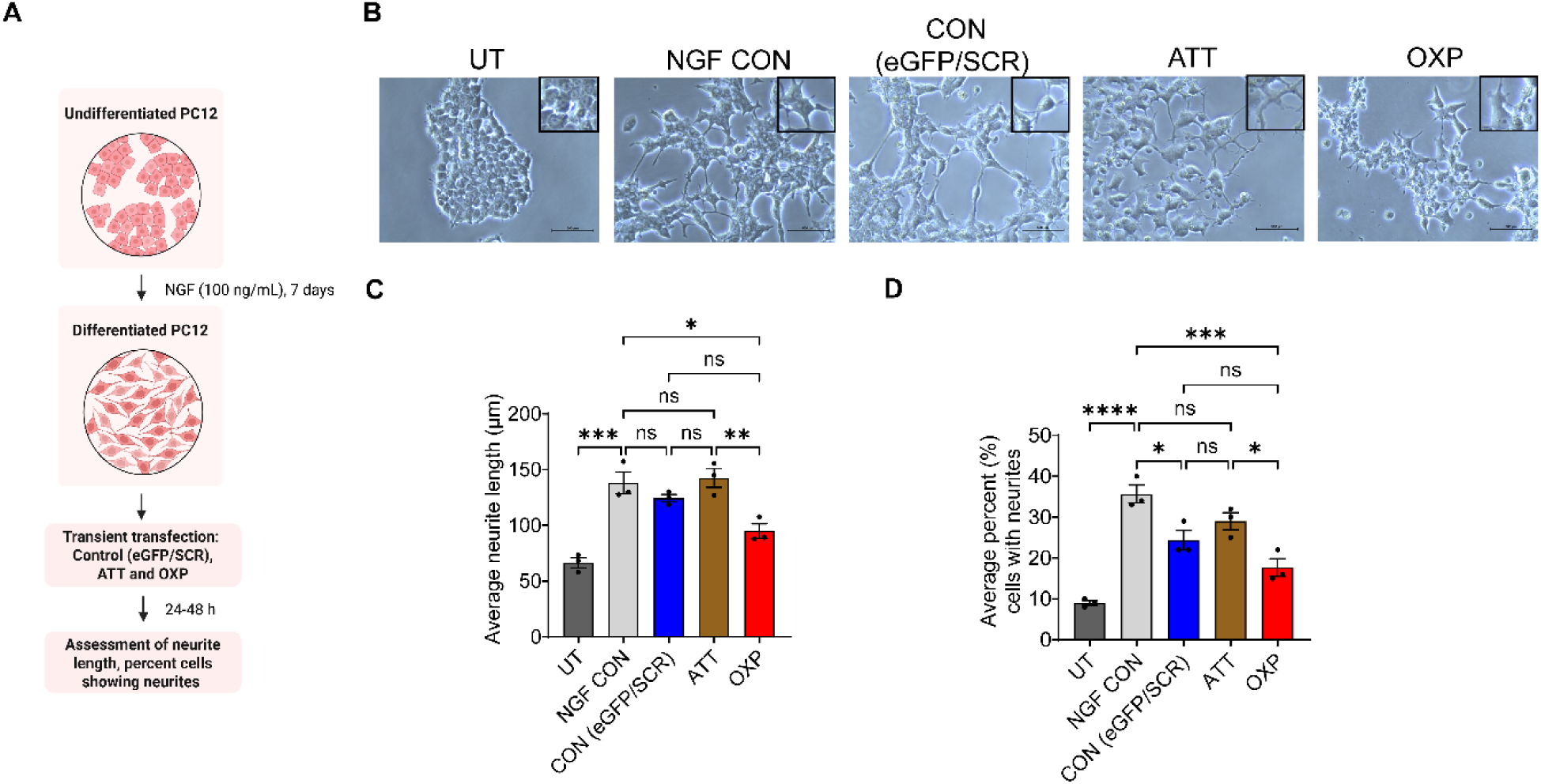
PLD1 Overexpression Reduces Neurite Number and Length in Differentiated PC12 Cells. (A) Schematic illustration of the PC12 cell differentiation and transfection process. PC12 cells were differentiated with NGF (100 ng/mL) for 7 days, followed by transient transfection. (B) Photomicrographs of PC12 cells treated with or without (UT) NGF (100 ng/mL) for 7 days, followed by transient transfection with CON (eGFP/SCR), ATT (PLD1 shRNA), and OXP (PLD1 OXP) plasmid constructs for 48 h in Opti-MEM. Scale bar = 500 μm. (C) Average neurite length (μm) in transfected PC12 cells. Bars represent mean ± SEM (N = 3 independent experiments with 6-7 images/treatment group). (D) Average percent (%) of cells with neurites in transfected PC12 cells. Bars represent mean ± SEM (N = 3 independent experiments with 6-7 images/treatment group). Statistical significance: ns = not significant, *p < 0.05; **p < 0.01, ***p < 0.001, multiple comparisons.

Our results demonstrate that PLD1 overexpression (OXP) significantly reduced both the average neurite length (Fig. 1C) and the average percentage of cells bearing neurites (Fig. 1D) in differentiated PC12 cells. This finding suggests that elevated PLD1 activity can impair neurite outgrowth, a process critical for establishing and maintaining synaptic connections, when administered post-differentiation. Conversely, while PLD1 attenuation (ATT) did not significantly increase neurite outgrowth beyond the control, it also did not inhibit it, indicating that reducing PLD1 levels is not detrimental.

### AAV2-Mediated Modulation of PLD1 Expression in the Hippocampus

Adeno-Associated Virus (AAV) vectors have emerged as powerful and widely adopted tools for gene delivery and manipulation within the CNS, making them ideal for our investigation (Weinberg et al. 2013). AAVs are ideally suited for efficient transduction in non-dividing, post-mitotic mature neuronal tissues thus targeting the adult and aging brain with lasting transgene expression primarily through the formation of stable extrachromosomal episomes. Compared to other viral vector systems, AAVs generally exhibit lower immunogenicity and pathogenicity, minimizing confounding inflammatory responses and cellular toxicity.

To confirm the successful transduction of our AAV2 vectors and visualize the resulting changes in PLD1 expression, we performed immunohistochemical analyses in aged WT and 3xTg-AD mice (Fig. 2). First, we used intra-hippocampal (CA1) stereotaxic surgery to establish specific expression patterns. We injected 3 µL of the control vector intracerebroventricularly and observed strong expression patterns for eGFP. We also injected PLD1 OXP and confirmed increased expression of PLD1, without eGFP expression since that GFP sequence was removed to insert the hPLD1 cDNA sequence (see Suppl. Fig. S2). In the VECTOR eGFP control group, we observed widespread eGFP expressions (green), indicating successful viral transduction, and endogenous PLD1 staining (red), showing the baseline distribution of PLD1 in the hippocampus. In the PLD1 OXP group, we observed a clear increase in PLD1 staining (red) compared to the control. This confirms that AAV2-mediated delivery of the PLD1 OXP construct effectively increased PLD1 levels in the transduced cells. Conversely, in the PLD1 ATT group, PLD1 staining (red) was markedly reduced in the hippocampal regions expressing eGFP (green), demonstrating the efficacy of the shRNA in attenuating PLD1 expression.

**Figure 2:**
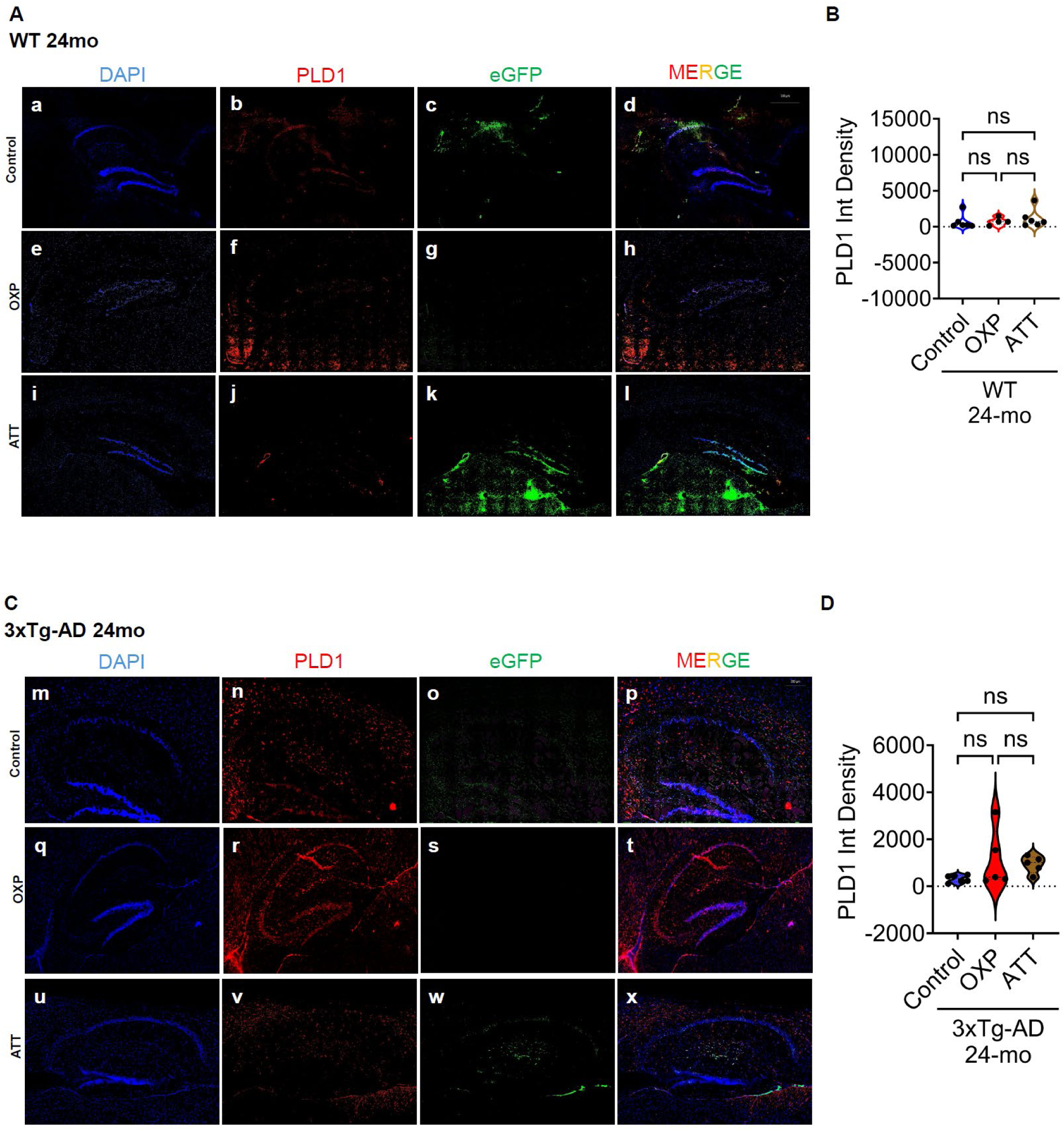
AAV2-mediated modulation of PLD1 expression in the hippocampus of 24-month-old Wild-Type (WT) and 3xTg-AD mice. Mice received hippocampal injections of AAV2 vectors at 18 months of age, designed to either overexpress human PLD1 (PLD1 OXP) or attenuate endogenous PLD1 using an shRNA (PLD1 ATT, co-expressing eGFP). Brain tissue was collected at 24 months for immunofluorescence analysis following behavioral and electrophysiological studies. Representative images show DAPI (blue, nuclei), PLD1 (red), and eGFP (green) staining, and merged channels in (A) WT and (C) 3xTg-AD mice. (a-d) WT mice injected with a control AAV2 vector (expressing eGFP) show baseline PLD1 expression and eGFP signal in the hippocampus. (e-h) WT mice were injected with AAV2-PLD1 OXP. These images display PLD1 levels in WT mice following attempted overexpression (PLD1 OXP construct does not contain an eGFP tag). (i-l) WT mice injected with AAV2-PLD1 ATT (shRNA-eGFP). Shows significant reduction in PLD1 immunofluorescence (j) in eGFP-positive cells (k), indicating successful shRNA-mediated knockdown of PLD1 (l, merge). (m-p) 3xTg-AD mice injected with a control AAV2 vector (expressing eGFP) show baseline PLD1 expression and eGFP signal in the hippocampus. (q-t) 3xTg-AD mice injected with AAV2-PLD1 OXP. (u-x) 3xTg-AD mice injected with AAV2-PLD1 ATT (shRNA-eGFP). Similar to WT mice, these images demonstrate a marked decrease in PLD1 protein levels (r) in hippocampal cells expressing eGFP (s), confirming effective PLD1 attenuation (t, merge). Scale bars (500 μm) are indicated on the merged images. Violin plots represent mean ± SEM (N=3 mice). Statistical significance: ns = not significant. Each circle represents one mouse. Data is represented as mean ± SEM. These results confirm the targeted modulation of PLD1 expression in the hippocampus of aged WT and 3xTg-AD mice.

Further such immunohistochemical findings also validate the ability of our AAV2 vectors to effectively modulate PLD1 expression in the hippocampus of aged WT and 3xTg-AD mice (Fig. 2). The observed increase in PLD1 expression with PLD1 OXP and the reduction with PLD1 ATT provide critical evidence for the specificity and efficacy of our gene transfer approach. These results, combined with our *in vitro* validation of the plasmid constructs (as demonstrated in Suppl. Fig. S2), confirm that we successfully achieved targeted modulation of PLD1 levels in our *in vivo* experiments.

To assess the validity of the AAV2 vectors in other brain regions, we performed immunohistochemical analysis of PLD1 expression in the frontal cortex region of aged WT (Suppl Fig. S11A-B) and 3xTg-AD mice (Suppl Fig. S11C-D). In both WT and 3xTg-AD cohorts, we observed a weak expression of PLD1 in the vector control group, while there was no significant change in PLD1 expression in the PLD1 ATT (ATT) and PLD1 OXP (OXP) group respectively.

### Attenuating PLD1 Promotes Improved Cognition in Aged WT and 3xTg-AD Mice

The core hypotheses regarding PLD1 in aging and synaptic vulnerability were tested in vivo using established mouse models. Using C57BL/6 or 129/SvJ mice aged to 20-24 months we assessed normal, non-pathological aging vs 3xTg-AD (age-matched) with advanced pathology to dissect the contributions of PLD1 to synaptic vulnerability arising from "normal" aging processes versus those specifically driven or exacerbated by AD/ADRD pathology thus providing a nuanced understanding of the role of PLD1 in synapses.

To investigate the effects of PLD1 OXP and PLD1 ATT on cognitive function, we performed a behavioral battery of tests at 20-months and again at 24-months, following a 4-month interval (to assess longitudinal effects) in the home cage (see Graphical Abstract). Y-maze, NOR, and FC tests, each assessed distinct aspects of memory and cognition.

The Y-maze test, which measures spontaneous alternation behavior, was used to assess spatial working memory (Bryan et al. 2009). Attenuating PLD1 expression significantly improved spontaneous alternation in 20-month-old 3xTg-AD mice (Fig. 3A), indicating enhanced working memory. Specifically, PLD1 ATT resulted in a 170.2% increase in spontaneous alternation compared to the control group (Con 100.0 ± 4.72 vs ATT 170.2 ± 18.99, ****p < 0.0001). While PLD1 OXP treatment also showed a trend of increase (18.4%) in 3xTg-AD mice (Con 100.0 ± 4.72 vs OXP 118.4 ± 9.90, ns), this change was not statistically significant. In contrast, neither PLD1 ATT (8.8% increase, Con 100.0 ± 6.38 vs ATT 108.8 ± 7.45, ns) nor PLD1 OXP (13.4% increase, Con 100.1 ± 6.38 vs OXP 113.4 ± 7.55, ns) significantly altered spontaneous alternation in 20-month-old WT mice (Fig. 3A).

**Figure 3:**
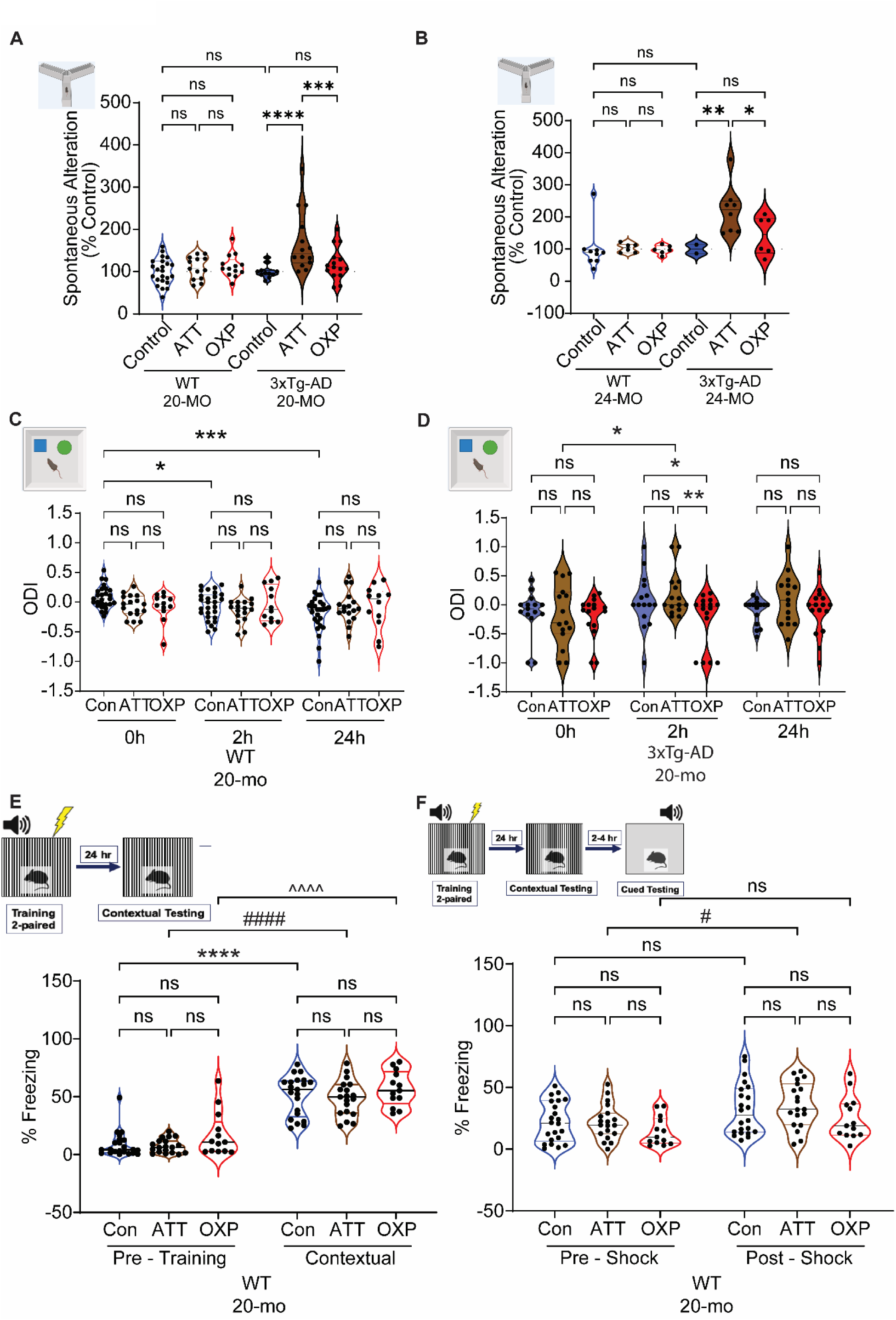
Attenuation of PLD1 Mitigates Cognitive Impairment in Aged WT and 3xTg-AD Mice. (A) Schematic timeline of AAV2 (eGFP, OXP, SCR, and ATT) administration followed by memory and cognitive behavior assessment in 20-month-old WT and 3xTg-AD mice. (B) Percentage of spontaneous alteration in 20-month-old mice. Violin plots depict the distribution of data points. Hollow violin represents WT, filled violin represents 3xTg-AD. Violin plots represent mean ± SEM (N=5-16 mice). (C) Percentage of spontaneous alteration in 24-month-old mice. Violin plots depict the distribution of data points. Hollow violin represents WT, filled violin represents 3xTg-AD. Bars represent mean ± SEM (N=5-16 mice). (D) Object Discrimination Index (ODI) in 20-month-old WT mice at 0 hours, 2 hours, and 24 hours. Violin plots depict the distribution of data points. Bars represent mean ± SEM (N=5-16 mice). (E) Object Discrimination Index (ODI) in 24-month-old 3xTg-AD mice at 0 hours, 2 hours, and 24 hours. Violin plots depict the distribution of data points. Bars represent mean ± SEM (N=5-16 mice). (F) Percentage of freezing in 20-month-old WT mice during pre-training, training, and contextual testing phases of fear conditioning. Violin plots depict the distribution of data points. Bars represent mean ± SEM (N=5-16 mice). (G) Percentage of freezing in 20-month-old WT mice during pre-shock, and post-shock phases of fear conditioning. Violin plots depict the distribution of data points. Bars represent mean ± SEM (N=5-16 mice). Statistical significance: ns = not significant, *p < 0.05, **p < 0.01, ****p < 0.0001. Each circle represents one mouse. Data are represented as mean ± SEM.

When mice were re-tested at 24 months, PLD1 attenuation continued to show a beneficial effect in 3xTg-AD mice. PLD1 ATT treatment resulted in a significant 222% increase in spontaneous alternation (Con 100.0 ± 13.80 vs ATT 222.0 ± 26.85, **p < 0.01) while PLD1 OXP treatment showed only a non-significant 41.9% increase (Con 100.0 ± 13.54 vs OXP 141.9 ± 24.76, ns). In aged WT mice, neither PLD1 ATT (3.5% increase, Con 100.0 ± 22.51 vs ATT 103.5 ± 5.76, ns) nor PLD1 OXP (2.34% decrease, Con 100.0 ± 22.52 vs OXP 97.66 ± 5.52, ns) significantly affected spontaneous alternation. These results suggest that PLD1 attenuation improves working memory in 3xTg-AD mice longitudinally over 4 months (Fig. 3B).

To assess hippocampal-dependent memory, we performed NOR (Krishnan et al. 2018; Bourne et al. 2019; Natarajan et al. 2023). In 20-month-old WT mice, neither PLD1 ATT nor PLD1 OXP treatment significantly affected the object discrimination index (ODI) at 0, 2, or 24 hours (Fig. 3C). This indicates that PLD1 modulation did not significantly impair object recognition memory in 20-month-old WT mice. In 20-month-old 3xTg-AD mice, however, we observed some trends, although most did not reach statistical significance (Fig. 3D). At 0 hours (training), both PLD1 ATT (34.2% decrease, Con −0.1376 ± 0.1005 vs ATT −0.1847 ± 0.1360, ns) and PLD1 OXP (41.5% decrease, Con −0.1376 ± 0.1005 vs OXP −0.1947 ± 0.09015, ns) showed non-significant trends towards decreased ODI. At 2 hours, PLD1 ATT showed a non-significant trend towards increased ODI (Con 0.05672 ± 0.1213 vs ATT 0.1780 ± 0.09898, ns), while PLD1 OXP significantly decreased ODI (538.9%, Con 0.05672 ± 0.1213 vs OXP −0.249 ± 0.1149, *p < 0.05). At 24 hours, both PLD1 ATT (611.9% increase, Con −0.0112 ± 0.05734 vs ATT 0.08812 ± 0.1103, ns) and PLD1 OXP (902.6% decrease, Con −0.0112 ± 0.05734 vs OXP −0.1123 ± 0.09737, ns) showed non-significant trends towards increased and decreased ODI, respectively. These results suggest that PLD1 overexpression may transiently disrupt object recognition memory in 3xTg-AD mice at the 2-hour time point, while PLD1 ATT did not have a significant impact.

FC was used to assess both hippocampal (contextual) and amygdala (cued) dependent associative memory in 20-month-old WT mice (Bourne et al. 2019; Natarajan et al. 2023). During the pre-training phase, when mice were introduced to the chamber, there were no significant differences in the percentage of freezing behavior among the control, ATT, and OXP groups (Con 8.513 ± 2.343 vs ATT 7.405 ± 1.387 and OXP 17.44 ± 5.354, ns), indicating similar baseline anxiety levels (Fig. 3E). However, during the contextual testing phase, when mice were returned to the chamber, all groups showed a significant increase in freezing compared to their pre-training levels, demonstrating successful learning (Con pre-training 8.513 ± 2.343 vs Con contextual 51.13 ± 3.763, ****p < 0.0001; ATT pre-training 7.405 ± 1.387 vs ATT contextual 48.78 ± 3.496, _####_p < 0.0001; OXP pre-training 17.44 ± 5.354 vs OXP contextual 56.99 ± 4.208, ^^^^p < 0.0001). There were no significant differences in contextual freezing among the treatment groups.

In the cued testing phase, where a tone was presented, there were no significant differences in freezing behavior among the groups before the shock (Pre-shock: Con 21.99 ± 3.470 vs ATT 21.84 ± 3.221 and OXP 14.57 ± 3.251, ns) or after the shock (Post-shock: Con 32.07 ± 4.462 vs ATT 35.95 ± 4.209 and OXP 25.11 ± 4.912, ns). However, PLD1 ATT significantly increased freezing from the pre-shock to the post-shock phase (Pre-shock: ATT 21.84 ± 3.221 vs Post-shock: ATT 35.95 ± 4.209, *p < 0.05), suggesting improved cued memory recall. PLD1 OXP showed a similar trend, but it did not reach statistical significance (Pre-shock: OXP 14.57 ± 3.251 vs Post-shock: OXP 25.11 ± 4.912, ns) (Fig. 3F).

In summary, our behavioral analyses demonstrate that PLD1 ATT improves spatial working memory in 3xTg-AD mice and enhances cued memory recall in WT mice. PLD1 OXP administration generally did not significantly impair cognitive function in these aged models. Given that anxiety-like behaviors are also affected in both healthy and pathological aging and considering our own unpublished findings indicating a role for PLD1 in anxiety regulation, we next investigated the effects of PLD1 modulation on these behaviors.

### Attenuation of PLD1 Modulates Anxiety- and Depression-like Behavior in Aged WT and 3xTg-AD Mice

To assess the impact of PLD1 modulation on anxiety-like behavior, we conducted elevated plus maze (EPM) tests in aged WT and 3xTg-AD mice. In 20-month-old WT mice, PLD1 ATT showed a trend towards decreasing the percentage of time spent in the open arms zone by 51.02% compared to the control cohort (Con 100 ± 29.78 vs ATT 48.98 ± 16.27, ns). Conversely, PLD1 OXP showed a trend towards increasing the percentage of time spent in the open arms zone by 31.7% compared to the control cohort (Con 100 ± 29.78 vs OXP 131.7 ± 50.58, ns), although neither trend reached statistical significance (Fig. 4A).

**Figure 4:**
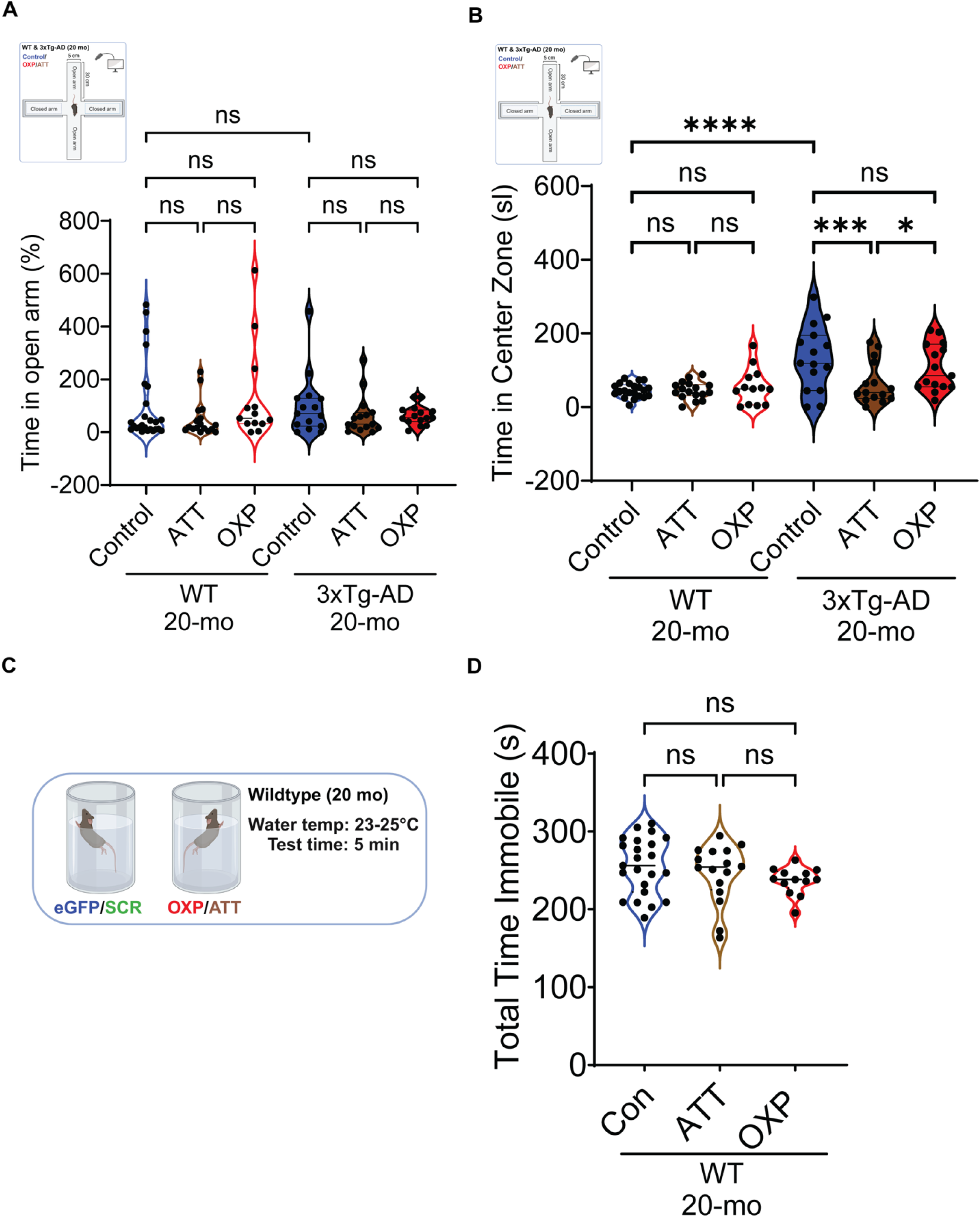
Attenuation of PLD1 Reduces Anxiety- and Depression-like Behaviors in Aged WT and 3xTg-AD Mice. Elevated plus maze (EPM) for control (SCR and eGFP combined), OXP, and ATT cohorts of 20-month-old WT and 20-month-old 3xTg-AD mice. (A) Time spent in the open arms (% control) in the EPM for OXP and ATT cohorts compared to the control cohort in 20-month-old WT and 20-month-old 3xTg-AD mice. Violin plots depict the distribution of data points. Data is represented as mean ± SEM (N=5-16 mice). Hollow violin plots represent WT, filled violin plots represent 3xTg-AD. (B) Time spent in the center zone (seconds) in the EPM for OXP and ATT cohorts compared to the control cohort in 20-month-old WT and 20-month-old 3xTg-AD mice. Violin plots depict the distribution of data points. Data is represented as mean ± SEM (N=5-16 mice). (C) Pictorial representation of the forced swim test (FST) for the 20-month-old WT mice. (D) Total time immobile (seconds) in the forced swim test (FST) for OXP and ATT cohorts compared to the control cohort in 20-month-old WT mice. Violin plots depict the distribution of data points. Data is represented as mean ± SEM (N=5-16 mice). Statistical significance: ns = not significant, *p < 0.05, ***p < 0.001, ****p < 0.0001. Each circle represents one mouse. Data are represented as mean ± SEM.

In 20-month-old 3xTg-AD mice, both PLD1 ATT and PLD1 OXP treatment groups showed a trend towards decreasing the percentage of time spent in the open arms zone compared to the control group (Con 100.0 ± 30.10 vs ATT 62.43 ± 19.36, ns; Con 100.0 ± 30.10 vs OXP 63.52 ± 9.453, ns). The ATT group showed a 37.57% decrease, and the OXP group a 36.48% decrease; however, neither change was statistically significant (Fig. 4A).

Next, we examined time spent in the center zone of the EPM, another measure often associated with anxiety-like behavior. In 20-month-old WT mice, PLD1 ATT did not significantly alter the time spent in the center zone compared to the control cohort (Con 45.39 ± 4.218 vs ATT 45.06 ± 5.744, ns). PLD1 OXP, however, showed a non-significant trend towards a 22.03% increase in time spent in the center zone (Con 45.39 ± 4.218 vs OXP 55.39 ± 13.78, ns) (Fig. 4B).

In 20-month-old 3xTg-AD mice, PLD1 ATT) resulted in a significant 50.81% decrease in time spent in the center zone (Con 132.2 ± 22.71 vs ATT 65.02 ± 14.82, ***p < 0.001). PLD1 OXP showed a non-significant trend towards a 19.36% decrease (Con 132.2 ± 22.71 vs OXP 106.6 ± 16.35, ns). Interestingly, the PLD1 OXP group spent significantly more time in the center zone compared to the PLD1 ATT group in 3xTg-AD mice (OXP 106.6 ± 16.35 vs ATT 65.02 ± 14.82, *p < 0.05). Furthermore, the 3xTg-AD control group displayed a significantly higher percentage of time spent in the center zone (191.25%) compared to the age-matched WT control group (3xTg-AD Con 132.2 ± 22.71 vs WT Con 45.39 ± 4.218, ****p < 0.0001), indicating an increased anxiety-like phenotype in the 3xTg-AD mice (Fig. 4B).

In summary, our EPM results suggest that PLD1 modulation influences anxiety-like behavior. PLD1 ATT tends to reduce anxiety-like behavior, particularly in 3xTg-AD mice, as indicated by decreased time in the center zone. PLD1 OXP, on the other hand, showed a trend towards increased anxiety-like behavior in WT mice, while its effects in 3xTg-AD mice were less pronounced.

To further investigate the effect of PLD1 modulation on depression-like behavior, we performed the forced swim test (FST) in 20-month-old WT mice (Fig. 4C). PLD1 ATT showed a non-significant trend towards a 3.83% decrease in total immobility time compared to the control cohort (Con 255.8 ± 7.579 vs ATT 246.0 ± 9.394, ns). Similarly, PLD1 OXP showed a non-significant trend towards a 7.5% decrease in total immobility time compared to the control cohort (Con 255.8 ± 7.579 vs OXP 236.6 ± 4.852, ns) (Fig. 4D). These results suggest that PLD1 modulation, in this context, does not significantly alter depression-like behavior in aged WT mice.

Taken together, our behavioral findings indicate that PLD1 modulation influences both cognitive function and anxiety-like behavior. We observed that PLD1 ATT generally improves spatial working memory and reduces anxiety-like behavior, while PLD1 OXP impaired object recognition memory and increased anxiety-like behavior in aged mice.

### Modulating PLD1 Differentially Alters Potentiation Profiles in the Schaffer Collateral Synapses of Aged WT and 3xTg-AD Mice

To investigate the role of PLD1 in synaptic function during aging and in the context of AD/ADRD pathology, we performed ex vivo electrophysiological recordings in hippocampal slices from AAV administered 24-month-old WT and 3xTg-AD mice. We focused on assessing HFS-LTP in SC - CA3->CA1 synapse, a well-established measure of synaptic plasticity (Krishnan et al. 2018; Bourne et al. 2019; Natarajan et al. 2023). Electrophysiological assessments were conducted at 24 months to align with longitudinal behavioral analyses performed from 20 to 24 months, following ICV administration at 18 months in the same cohorts. Previous work from our group has demonstrated that reducing PLD1 activity pharmacologically can mitigate the detrimental effects of amyloidogenic proteins (AβO or TauO) on HFS-LTP (Krishnan et al. 2018), suggesting that PLD1 contributes to amyloid-mediated synaptic dysfunction. The current study extends these findings by directly manipulating PLD1 expressions to examine its intrinsic influence on synaptic vulnerability during aging.

In aged WT mice, PLD1 OXP significantly impaired LTP compared to controls (Fig. 5A and C, WT Con: 269.5 ± 7.405 vs WT OXP: 139.7 ± 7.985, ****p<0.0001). This result indicates that excessive PLD1 activity disrupts the ability of synapses to undergo long-term strengthening in otherwise healthy aged mice. Conversely, attenuation of PLD1 ATT did not significantly alter LTP compared to WT controls (Fig. 5A and C, WT Con: 269.5 ± 7.405 vs WT ATT: 259 ± 22.62, ns), suggesting that reducing PLD1 levels does not compromise synaptic plasticity in aged WT mice.

**Figure 5:**
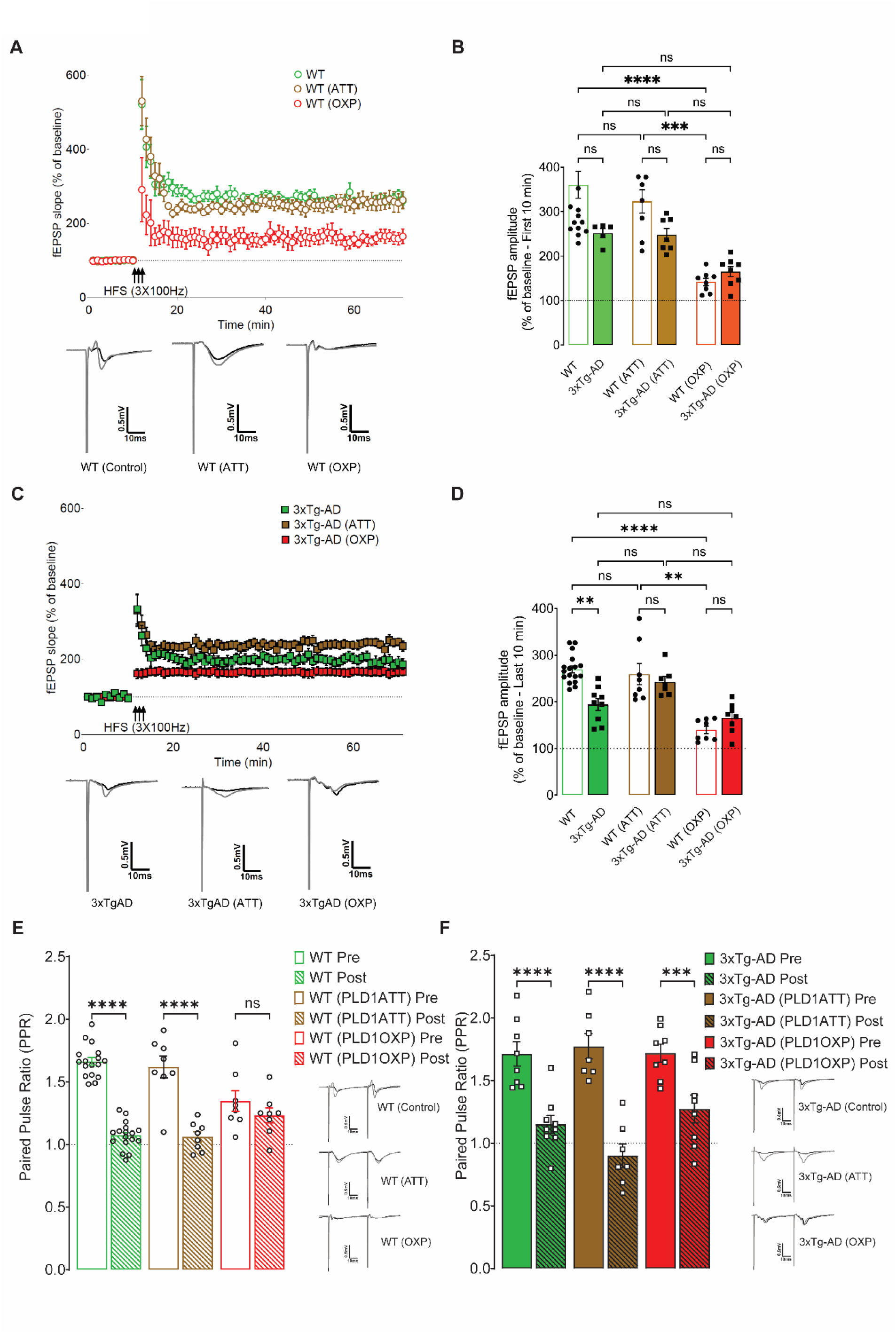
Attenuation of PLD1 Rescues Long-Term Potentiation and Paired-Pulse Ratio in Aged Wild-Type and 3xTg-AD Mice. (A) Time course of field excitatory postsynaptic potential (fEPSP) slope (% of baseline) following high-frequency stimulation (HFS, 3x100Hz) in 24-month-old wild-type (WT) mice. Green circles represent WT (control), brown circles represent WT treated with ATT, and red circles represent WT treated with OXP. Inset traces depict representative fEPSP recordings before (black) and after (grey) HFS. (B) Quantification of fEPSP slope (% of baseline) during the first 10 minutes post-HFS in WT (hollow bars) and 3xTg-AD (filled bars) mice respectively. Bars represent mean ± SEM (N=3-4 mice). (C) Time course of fEPSP slope (% of baseline) following HFS in 24-month-old 3xTg-AD mice. Green squares represent 3xTg-AD (control), brown squares represent 3xTg-AD treated with ATT, and red squares represent 3xTg-AD treated with OXP. Inset traces depict representative fEPSP recordings before (black) and after (grey) HFS. (D) Quantification of fEPSP slope (% of baseline) during the last 10 minutes post-HFS in WT (hollow bars) and 3xTg-AD (filled bars) mice respectively. Bars represent mean ± SEM (N=3-4 mice). (E) Paired-pulse ratio (PPR) in WT mice. Bars represent mean ± SEM (N=3-4 mice). Circles represent individual data points. Inset traces show representative fEPSP recordings for each condition. (F) Paired-pulse ratio (PPR) in 3xTg-AD mice. Bars represent mean ± SEM (N=3-4 mice). Squares represent individual data points. Inset traces show representative fEPSP recordings for each condition. Statistical significance: ns = not significant, **p < 0.01, ****p < 0.0001. Data is represented as mean ± SEM.

In the 3xTg-AD model of pathological aging, the effects of PLD1 modulation were distinct. While there was a trend towards improved synaptic potentiation with PLD1 ATT at the last 10 minutes of recording (Fig. 5D and F, 3xTg-AD: 194 ± 12.62 vs 3xTg-AD (ATT): 242.9 ± 11.43 vs 3xTg-AD (OXP): 165.7 ± 11.30, ns), this difference did not reach statistical significance. This suggests that reducing PLD1 may have a protective effect against synaptic dysfunction in this late-onset AD/ADRD model, though the effect is less pronounced than the detrimental impact of OXP in WT mice. Notably, PLD1 OXP in 3xTg-AD mice also showed a trend towards lowered potentiation compared to both controls and the ATT group, but again, this did not achieve statistical significance. This could imply that the 3xTg-AD model already exhibits a degree of synaptic compromise, potentially limiting the additional disruptive effect of further PLD1 OXP.

To further investigate the mechanisms underlying these changes in LTP, we examined post-tetanic potentiation (PTP), a measure of presynaptic release probability (Fig. 5B and Fig. 5E). In both WT and 3xTg-AD mice, PLD1 OXP was associated with a significant decrease or a trend of decrease in vesicular release probability. This observation aligns with our previous findings in amygdala-dependent memory recall studies, where elevated PLD1 impaired presynaptic function (Krishnan et al. 2016; Krishnan 2016; Krishnan et al. 2011). These findings suggest that excess PLD1 disrupts the ability of synapses to efficiently release neurotransmitter, regardless of the presence of AD-related pathology.

Paired-pulse facilitation (PPF), another measure of presynaptic function, revealed further insights (Fig. 5G – WT and H – 3xTg-AD). In WT mice, there was a significant difference between pre- and post-PPF values in the control [WT Con (Pre: 1.663 ± 0.03119 vs Post: 1.073 ± 0.02617, ****p<0.0001)] and ATT [WT PLD1 ATT (Pre: 1.619 ± 0.08711 vs Post: 1.061 ± 0.04040, ****p<0.0001)] groups, indicating normal presynaptic modulation.

However, this difference was absent in the WT PLD1 OXP group [WT OXP (Pre: 1.347 ± 0.08312 vs Post: 1.233 ± 0.05954, ns)], suggesting that PLD1 OXP impairs only the presynaptic mechanisms in aged WT mice.

Interestingly, this disruption of PPF by PLD1 OXP was not observed in the 3xTg-AD cohort (3xTg-AD Pre: 1.713 ± 0.09657 vs Post: 1.150 ± 0.07157, ****p<0.0001; 3xTg-AD PLD1 ATT Pre: 1.774 ± 0.1007 vs Post: 0.9007 ± 0.09502, ****p<0.0001; 3xTg-AD PLD1

OXP Pre: 1.720 ± 0.07165 vs Post: 1.273 ± 0.1110, ***p=0.0001). This apparent discrepancy could be attributed to compensatory mechanisms or ceiling effects in the 3xTg-AD model, where elevated basal PLD1 levels may already contribute to presynaptic dysfunction, limiting the additional impact of further overexpression. This idea aligns with our previous findings and clinical observations of elevated PLD1 in AD (Krishnan et al. 2018; McKenzie et al. 2017) but requires further mechanistic investigation.

To complement our LTP and PPF findings, we assessed input-output (I/O) relationships by measuring fEPSP slope as a function of fiber volley (FV) amplitude (Suppl. Fig. S5 for WT and S6 for 3xTg-AD). In WT mice, the ATT group displayed I/O profiles similar to controls (Suppl. Fig. S5 D, E, F compared to A, B, C respectively), further supporting the notion that reducing PLD1 does not impair basal synaptic transmission in aged WT animals. However, PLD1 OXP resulted in a marked reduction in fEPSP slope as a function of fiber volley (Suppl. Fig. S5 G, H, I compared to A, B, C respectively), indicating impaired synaptic efficacy due to PLD1 upregulation in aging WT mice.

In 3xTg-AD mice, PLD1 ATT enhanced the post-HFS I/O slope compared to controls (Suppl. Fig. S6 D, E, F compared to A, B, C respectively), suggesting a potential rescue of synaptic function. Conversely, PLD1 OXP significantly disrupted both pre- and post-HFS I/O profiles in 3xTg-AD mice (Suppl. Fig. S6 G, H, I compared to A, B, C respectively), proving the hypothesis that elevated PLD1 contributes to synaptic vulnerability in this late-onset AD/ADRD model.

Cumulative probability analyses of LTP (Suppl. Fig. S7 A, B) and PTP (Suppl. Fig. S7 C, D) further illustrate the differential effects of PLD1 modulation. These analyses reveal that OXP shifts the frequency distribution of synaptic potentiation towards lower values (leftward shift) compared to control, irrespective of whether it is WT or 3xTg-AD.

To further investigate the spatial and temporal dynamics of synaptic plasticity, we utilized multi-electrode arrays (MEAs) (Liu et al. 2012). MEA technology allows for simultaneous recording of electrical activity (spikes and local field potentials, LFPs) from multiple sites across the hippocampal slice. This provides a powerful means to investigate, not just single-synapse plasticity, but also emergent network properties, such as spontaneous firing patterns, network oscillations, synchrony between neuronal populations, and the propagation of evoked activity across the hippocampal circuit.

MEA recordings in hippocampal slices from aged WT and 3xTg-AD mice corroborated and expanded the findings from conventional electrophysiology (Suppl. Fig. S8-10). Qualitative LTP heatmaps revealed an overall attenuation of response amplitude in 3xTg-AD mice compared to WT (Suppl. Fig. S8A and B). In WT slices, PLD1 ATT resulted in decreased response amplitude and an increased number of non-responsive electrodes, while PLD1 OXP further attenuated response amplitude and, surprisingly, induced a polarity flip in some areas. Analysis of electrode response types (depolarization, hyperpolarization, both, or none) revealed a trend towards an increased number of non-responsive areas with PLD1 modulation (Suppl. Fig. S9). PPF was reduced in WT mice with PLD1 ATT (Suppl. Fig. S10A), and while IO maximum amplitude was not significantly altered, AAV-mediated PLD1 modulation reduced variability compared to aged WT controls (Suppl. Fig. S10B). These MEA findings highlight the complex spatial and temporal alterations in synaptic activity induced by PLD1 modulation, suggesting that PLD1 not only influences synaptic strength but also network excitability and response patterns.

These findings support the hypothesis that elevated PLD1 contributes to synaptic vulnerability, whereas ATT appears to preserve or shift synaptic parameters toward higher values, suggesting a protective effect.

## DISCUSSION

Our multimodal assessments employing in vitro studies in differentiated PC12 cells and in vivo AAV2-mediated gene transfer in aged wild-type and 3xTg-AD mice, provide insights into how PLD1 specific modulation affects synaptic function in healthy and pathological states. In PC12 cells, PLD1 OXP impaired neurite outgrowth, while PLD1 ATT showed no detrimental effects. Using immunofluorescence, we validated levels of PLD1 modulation using our AAV2 system in the hippocampus and demonstrated that PLD1 ATT improved spatial working memory in 3xTg-AD mice and reduced anxiety-like behavior, while PLD1 OXP further lowered 3xTg-AD short-term memory in NOR compared to control 3xTg-AD group. Interestingly, altering PLD1 levels in wild-type mice did not result in any notable changes in cognitive or anxiety behavior. However, electrophysiological studies revealed that PLD1 OXP significantly impaired hippocampal HFS-LTP in wild-type mice and disrupted presynaptic function in both genotypes.

PLD1 exhibits dynamic subcellular localization. In resting cells, it is typically found in the perinuclear region, Golgi complex, and endosomes. However, upon cellular stimulation, PLD1 is translocated to the plasma membrane (Park & Han 2018). Crucially for neuronal function, PLD1 has been detected in synaptosomal preparations associated with the plasma membrane fraction (Humeau et al. 2001) that can impinge on membrane vesicle trafficking, cytoskeletal organization, cell proliferation and survival, cellular differentiation, signal transduction, exocytosis, and endocytosis. However, PLD1 presents a central paradox. Studies across multiple species have shown that knocking out the gene encoding PLD1 is not detrimental to survival (Panda et al. 2018; Bravo et al. 2018; Morris 2019). This observation suggests an existence of functional redundancy, perhaps via PLD2 or other compensatory lipid metabolic pathways, for its essential basal functions. Yet, despite this apparent dispensability for survival, a large body of evidence implicates the dysregulation of PLD1 – involving aberrant increases in its activity or expression – in the pathogenesis of neurodysfunctional states including AD (Jin et al. 2007; Jin et al. 2006; Kanfer et al. 1996; Krishnan et al. 2018), amyotrophic lateral sclerosis (ALS) (Kankel et al. 2020; May-Dracka et al. 2022), neuroimmune conditions like multiple sclerosis (MS) (Eftekharian et al. 2017; Göbel et al. 2014), and acute conditions like spinal cord injury (SCI) (Ke et al. 2024). We showed that PLD1 isoform expression and activity are elevated in crude neurosynaptosomal fractions from postmortem AD hippocampi (Krishnan et al. 2018; Bourne et al. 2019; Natarajan et al. 2023; Currie et al. 2024). Importantly, this elevated activity is causally linked to synaptic vulnerability and dysfunction. PLD1 acts as a key downstream mediator for the synaptotoxic effects of both AβO and TauO, and its inhibition confers significant protection against synaptic deficits and cognitive decline in preclinical models (Krishnan et al. 2018; Bourne et al. 2019; Natarajan et al. 2023; Currie et al. 2024). Comprehending the dichotomous roles of PLD1 – a critical developmental component and physiological regulator in mature systems, versus a potential instigator of pathology in disease states – is fundamental for elucidating basic synaptic mechanisms. This understanding is equally vital for the rational design of targeted, efficacious, and safe therapeutic interventions to preserve synaptic integrity and cognitive function against neurodegenerative conditions.

The inhibitory effect of PLD1 OXP on branching in differentiated cells (Fig. 1) stands in contrast to the established requirement for PLD1 during the initial outgrowth phase (Yoon et al. 2005). This suggests that the observed phenotype is likely a consequence of a "gain-of-function" effect, where exceeding the optimal physiological range of PLD1 activity becomes detrimental. In the context of a relatively mature neurite structure at Day 5, artificially high PLD1 levels appear to disrupt cellular homeostasis, possibly by over-activating intrinsic inhibitory mechanisms like the RhoA pathway or by causing imbalances in cytoskeletal regulation and membrane trafficking.

The finding that reducing PA levels can alleviate the branching inhibition caused by PLD1 OXP (Zhu et al. 2012) strongly points to PA as the key downstream molecule mediating this negative effect. The excess PA, generated by the overexpressed enzyme, likely interacts with downstream effectors that ultimately suppress the morphological changes associated with branching.

The second key observation is that partial reduction of PLD1 expression via shRNA in PC12 cells, differentiated with NGF for 5 days, did not lead to a significant change in neurite branching morphology compared to control cells (Fig. 1). The critical requirement for PLD1 activity appears most pronounced during the dynamic phase of neurite initiation and active elongation (Yoon et al. 2005). Once neurites are established and differentiation has reached a plateau (around Day 5-7) in PC12 cells (Das et al. 2004), the dependence on high levels of PLD1 activity for maintaining the gross structure might diminish. Mammalian PLD1 typically exhibits low basal activity compared to PLD2 (Park & Han 2018). It is plausible that the remaining PLD1 activity after partial shRNA knockdown, possibly supplemented by basal PLD2 activity, is sufficient to fulfill the maintenance requirements for the existing branching pattern. The primary role of PLD1 in more mature neuronal structures, therefore, might shift towards regulating synaptic functions, such as neurotransmitter release (Vitale et al. 2001) or dendritic spine dynamics (Luo et al. 2017), rather than maintaining the overall architecture of major branches. Alternatively, RhoA, a small GTPase known to regulate cytoskeletal dynamics and often associated with growth inhibition or retraction (Pearn et al. 2018), activates PLD1 (Zhu et al. 2012). The inhibition of branching caused by constitutively active RhoA can be partially rescued by knocking down PLD1 (Zhu et al. 2012). Furthermore, the enzymatic product of PLD1, PA, appears to be a key mediator, as reducing PA levels can partially ameliorate the branching inhibition caused by active RhoA or PLD1 overexpression (Zhu et al. 2012).

A side-by-side comparison of cognitive performance in 3xTg-AD and WT mice at advanced ages (20-24 months) reveals the specific impact of AD/ADRD-like pathology superimposed upon the background of normal aging. Utilizing a genetic approach with AAV2 vectors to either overexpress (PLD1 OXP) or attenuate (PLD1 ATT) PLD1 levels in aged (18-month-old at injection) wild-type (WT) and 3xTg-AD mice, we conducted longitudinal behavioral assessments (at 20 and 24 months) and terminal electrophysiological analyses (at 24 months) and focused it on how PLD1 signaling impinges under these different circumstances. PLD1 OXP exacerbates hippocampal synaptic dysfunction, particularly in aged WT mice, and impairs specific cognitive functions while PLD1 ATT appears protective, reducing synaptic vulnerability and improving spatial working memory and anxiety-like behaviors, most notably in the 3xTg-AD model. Thus, excessive signaling through the RhoA-PLD1-PA pathway, triggered by PLD1 OXP, could disrupt the delicate cytoskeletal balance needed for branching. This might manifest as excessive actin stabilization, increased contractility preventing protrusion, or impaired coordination between actin-based exploration and microtubule invasion required for branch consolidation, ultimately hindering branch formation or maintenance. While NGF itself promotes microtubule assembly essential for outgrowth (Drubin et al. 1985), supra-physiological levels of PLD1 activity might interfere with this process or its coordination with actin dynamics at potential branch points. On the other hand, the contrast between the detrimental effect of PLD1 OXP and the lack of effect upon partial attenuation points towards a non-linear relationship and a potential threshold effect. The differentiated neurite structure at Day 5 appears resilient to a moderate decrease in PLD1 activity, suggesting that the system can tolerate levels below the normal physiological range without immediate morphological consequence. This implies that while some basal PLD1 activity might be necessary, the cell has mechanisms to cope with a partial reduction but lacks mechanisms to effectively buffer the consequences of super physiological activity, particularly the overactivation of inhibitory pathways like RhoA.

In our previous studies, we employed pharmacological approaches to elucidate the pivotal role of attenuating aberrant PLD1 signaling in Aβ and tau-induced synaptic dysfunction and the underlying cognitive decline associated with AD/ADRD) in the 3xTg-AD mouse model (Krishnan et al. 2018; Bourne et al. 2019; Natarajan et al. 2023; Currie et al. 2024). Our findings demonstrated the preservation of dendritic spine integrity, thereby validating our current study conducted with PC12 cells. In this study, we directly validated, for the first time, PLD1 gene expression modulation in neurite changes, as depicted in Fig. 1. We, therefore, pursued AAV vector based post developmental modulations of the PLD1 gene in our aged mouse models (Fig. 3 through Fig. 5). The cognitive benefits observed with PLD1 attenuation in 3xTg-AD stand in stark contrast to the apparent lack of significant effects in healthy, aging WT animals under similar conditions (Fig. 3-5 and Suppl. Fig. S3). While acute PLD1 inhibition can effectively block the detrimental effects of exogenously applied AβO or TauO on synaptic function and memory processes in WT mice (Krishnan et al., 2018), the collective evidence strongly supports that chronic inhibition of PLD1 does not impinge the baseline cognitive performance in WT mice undergoing normal aging (Suppl. Fig. S1). These findings support a beneficial role for reduced post developmental PLD1 activity and expression. This is also confirmed by studies that show this conserved protein isoform (from bacteria to humans) to be neither lethal nor reproductively disabled even in germline knockouts of multiple species (Panda et al. 2018; Morris 2019). Importantly, this is the first study that reports the behavioral outcomes in a comprehensive manner for post developmental PLD1 ATT being non-detrimental to cognitive abilities in aged WT mice. The observation that PLD1 ATT yields significant behavioral benefits primarily in AD models while having minimal impact on healthy WT controls points towards mechanisms that are intrinsically linked to the pathological state. Several interconnected factors likely contribute to this state-dependency. In addition to the most straightforward explanation for the differential effect, PLD1 inhibition acts to normalize a pathway that is specifically hyperactive in the disease state. The impact of PLD1 modulation appears critically dependent on the presence of the specific molecular drivers of AD pathology, namely toxic forms of Aβ and tau. The functional state of synapses is fundamentally altered leading to impaired LTP, enhanced LTD, or a general dampening of the capacity for synaptic plasticity (Cuestas Torres & Cardenas 2020) including threshold modulations that alter plasticity disrupting synaptic homeostasis – the ability of neurons and circuits to adjust their properties to maintain stable function despite perturbations (Navakkode & Kennedy 2024). Modulation of PLD1 activity could therefore have a significantly greater impact on AD/ADRD than in healthy aging. The differential homeostatic adaptations observed between excitatory and inhibitory neurons even in response to a single perturbation (Lezmy et al. 2020) highlight the complexity and cell-type specificity of these processes, and give credence to such a speculation with altered PLD1 that we have validated in our published pharmacological studies (Krishnan et al. 2018; Bourne et al. 2019; Natarajan et al. 2023; Currie et al. 2024). AD neuropathology does not affect the brain uniformly but targets specific neuronal populations and large-scale networks, such as the default mode network (DMN) and circuits within the hippocampus and associated cortical areas (Li et al. 2024). Thus, PLD1 modulation might exert more pronounced effects within these already vulnerable or compromised circuits compared to relatively healthy and resilient circuits in WT. The enrichment of PLD1 expression in vulnerable regions like the hippocampus supports the potential for region-specific impacts (McDermott et al. 2020). In conclusion, our preclinical evidence in the present study (in conjunction with our published studies) strengthens our hypothesis that PLD1 ATT exerts state-dependent effects, offering significant cognitive and synaptic benefits primarily in the context of AD/ADRD pathology with minimal impact on healthy controls.

3xTg-AD mice exhibited an atypical baseline phenotype compared to WT controls, spending significantly more time in the relatively exposed center zone of the maze, a behavior not typically associated with heightened anxiety in rodents. Within this specific context, PLD1 ATT produced a statistically significant reduction in the time spent in the center zone by the 3xTg-AD mice which we interpret as “anxiolytic” within the specific behavioral profile of this animal model. In contrast, PLD1 OXP showed only non-significant trends towards reducing center time in 3xTg-AD mice, while in WT mice, there was a non-significant trend towards potentially increasing open arm/center time (anxiolytic-like). The anxiolytic effect of PLD1 ATT was observed only in the 3xTg-AD mice, and therefore we speculate that reducing PLD1 levels—particularly when they may be pathologically elevated as part of AD/ADRD (Krishnan et al. 2018) —could help to normalize the function of anxiety-related neural circuits that are dysregulated (including anxiety, depression, apathy, and agitation) in a vast majority (up to 90%) of AD patients (Li et al. 2024). Anxiety regulation involves a network of brain regions, prominently including the amygdala, hippocampus, prefrontal cortex (PFC), and anterior cingulate cortex (ACC) that are also among the earliest and most significantly affected by AD neuropathology (including Aβ deposition, tau tangles, neurodegeneration, and altered connectivity). Therefore, confirming PLD1 modulation in the hippocampus formation (Fig. 2)—a region impacting cognition, anxiety, and stress regulation via amygdala/PFC connections (Zhang et al. 2024)—we propose a direct or indirect influence on these circuits, building upon our previous work (Krishnan 2016). Interpreting the reduction in center time caused by PLD1 ATT as purely "anxiolytic" requires more assessments. However, the crucial finding remains that the statistically significant change induced by PLD1 ATT is relative to the own control group of age-matched 3xTg-AD. This change represents a shift towards the behavioral pattern implicitly observed in WT mice (which typically spend less time in the center), suggesting a normalizing or ameliorating effect of PLD1 ATT within the specific context of 3xTg-AD behavioral abnormalities. This highlights the importance of focusing on the intervention-induced relative change within a given model system, acknowledging that the precise human correlation of the baseline behavior may be complex. Nevertheless, targeting PLD1 might offer a dual benefit, potentially ameliorating both cognitive and non-cognitive symptoms, such as anxiety, in AD/ADRD neuropathology.

Immunohistochemical analyses assessing the cell-type distribution of PLD1 revealed that PLD1 colocalizes with both neurons and astrocytes. In 24-month-old WT mice, PLD1 displayed differential colocalization with Nrx1β across hippocampal subregions, including CA1, CA2, and the dentate gyrus (DG). Although PLD1 expression was lower in the ATT group, we observed a significant increase in its colocalization with Nrx1β, whereas the OXP group showed no detectable colocalization between PLD1 and Nrx1β (Suppl. Fig. S12A, C). A similar expression pattern was observed in 24-month-old 3xTg-AD mice. Notably, in the CA3 region, PLD1-Nrx1β colocalization (in both OXP and ATT) was significantly reduced relative to control group, while no significant differences were detected in the CA1 or DG regions of the hippocampus (Suppl. Fig. S12B, D). Furthermore, in 24-month-old WT mice, the OXP group showed a modest increase in PLD1-GFAP colocalization, whereas the ATT group exhibited a significant disruption of this interaction specifically in the CA3 region, but not in CA1 or the DG (Suppl. Fig. S12E, G). Interestingly, in 24-month-old 3xTg-AD mice, PLD1-GFAP colocalization in ATT was significantly reduced in CA1, while there was no change observed in the CA3 and DG (Suppl. Fig. S12F, H). Notably, PLD1 colocalization with GFAP in OXP was significantly elevated in the DG, but not in the CA1 or CA3.

A significant finding emerged from the electrophysiological assessment of synaptic plasticity concerning the differential impact of PLD1 modulation on HFS-LTP across genotypes. In aged WT mice, PLD1 OXP led to a pronounced and statistically significant impairment of LTP induction and maintenance compared to control animals. In contrast, within the cohort of aged (24-month-old) 3xTg-AD mice, neither PLD1 OXP nor PLD1 ATT resulted in statistically significant changes in LTP magnitude or persistence, although non-significant trends were noted towards diminished potentiation with OXP and potentially improved maintenance with ATT (Fig. 5). The advanced age (24 months) of the mice, combined with the severe Aβ and tau neuropathology typical of the 3xTg-AD model at this late stage, likely results in substantial pre-existing synaptic compromise. We propose that this scenario creates a potential "ceiling effect," limiting the system’s capacity to exhibit further significant changes in LTP in response to PLD1 manipulation. The concept of a ceiling effect (or, perhaps more accurately termed as "floor effect" when considering deficits in potentiation) operates distinctly for overexpression vs attenuation scenarios. A possible explanation supported by the broader AD/ADRD research suggests that synaptic machinery essential for LTP induction and maintenance (involving processes like NMDA receptor activation, calcium influx, kinase activation, and AMPA receptor trafficking and insertion (Prieto et al. 2017)) may already be operating at a significantly diminished capacity due to chronic exposure to pathological insults like Aβ and tau. Consequently, while the additional burden imposed by AAV2-mediated PLD1 overexpression is likely detrimental, it may fail to push the already severely impaired system even lower on the measured LTP scale. Synaptic efficacy (or strength) may already be operating close to a minimum functional level, thus hindering the detection of further statistically significant decreases. Furthermore, given reports that endogenous PLD1 expression and activity are already aberrantly elevated in AD brains and specifically increase with age in the hippocampus of 3xTg-AD mice (Krishnan et al. 2018), the exogenous overexpression might encounter a saturation point. If the downstream pathways mediating PLD1 detrimental effects on CA1 LTP are already maximally perturbed by the endogenous pathological elevation, adding more PLD1 via AAV2 might yield diminishing returns in terms of measurable functional impairment. Conversely, the potential benefits of reducing PLD1 levels via ATT might be insufficient to elicit a statistically significant rescue of LTP when confronted with severe, established pathology, thus explaining the positive trends but not reaching significance. Although we observed a significant reduction in LTP, this change did not translate into most of the behavioral assays we conducted, except for the y-maze assay, which reflects a more passive or innate behavior in 20–24-month-old WT mice. We attribute this limitation primarily to technical factors, including the timing of AAV injections (around 18 months), which may have reduced the likelihood of detecting a more robust PLD1 effect beyond the y-maze. To address this, we are designing experiments in which PLD1 OXP constructs will be administered earlier in post-developmental stages starting at 3, 6, or 12 months, to determine whether progressively elevated levels of aberrant PLD1 overexpression leads to broader behavioral deficits across multiple assays. Perhaps, administering such shRNA earlier, around the development of the pathology (3 months in 3xTg-AD) could result in robust improvement. The results highlight an inherent limitation in utilizing very late-stage disease models for evaluating interventions aimed at functional rescue. While valuable for modeling advanced human pathology, such models may lack the necessary dynamic range or sensitivity to detect statistically significant improvements (or further declines) in functional measures like HFS-LTP. Given that PLD1 ATT significantly improved behavioral outcomes (Y-maze, EPM) in these aged 3xTg-AD mice, the concurrent lack of effect on CA1 LTP suggests this assay alone might be insufficient in late-stage disease, likely due to ceiling/floor limitations. This finding underscores the necessity of employing diverse outcome measures, such as behavioral and MEA recordings that can potentially detect changes even when advanced pathology limits the responsiveness of any single electrophysiological measure.

MEA experiments revealed that in aged WT mice, modulating PLD1 levels induced alterations in the spatial patterns of synaptic responses following HFS, when compared to control slices. Specifically, both PLD1 ATT and OXP resulted in an increased number of electrodes that became non-responsive to the stimulation protocol after HFS. Furthermore, PLD1 OXP uniquely caused polarity flips—a switch from the typically negative-going fEPSPs to positive-going potentials—at some recording sites. The finding that both increasing (OXP) and decreasing (ATT) PLD1 levels altered the spatial map of potentiation suggests that basal PLD1 activity is intrinsically involved in maintaining the normal pattern of network activation and synaptic modification in the aging hippocampus. PLD1 activity directly influences the lipid composition of cellular membranes, particularly through the generation of phosphatidic acid (PtdOH) (McDermott et al. 2020), and is deeply integrated into the regulation of membrane vesicle trafficking and cytoskeletal dynamics. Perturbations in PLD1 levels could therefore impact neuronal excitability via changes in membrane lipid composition or downstream signaling that alters the function or localization of ion channels, thereby modifying intrinsic neuronal excitability, or synaptic transmission via (a) vesicle trafficking affecting neurotransmitter release probability (presynaptic effects, consistent with observed changes in PTP and PPF, see also (Krishnan et al. 2011; Krishnan et al. 2016) or (b) influence the density, localization, or function of postsynaptic receptors (postsynaptic effects) or (c) via interaction with proteins involved in trafficking pathways, such as presenilin-1 (PS1) and Rab GTPases, providing molecular links to these processes (Zhang et al. 2011). Alternatively, these alterations in cytoskeletal dynamics could be potentially influenced by PLD1 signaling pathways involving proteins like cofilin (Kang & Woo), that could affect dendritic integration or axonal conduction properties. The induction of polarity flips specifically by PLD1 OXP suggests a more profound disruption of network function. This could potentially reflect shifts in the local balance between excitatory and inhibitory synaptic currents, aberrant electrical field propagation due to altered cellular current sources/sinks, or changes in network synchrony. Aberrant network synchrony and dynamics are indeed implicated in AD pathophysiology (Andrade-Talavera & Rodríguez-Moreno 2021). PLD1 modulation also reduced response variability across electrodes which we speculate to be an indication of constriction in the network dynamic range or perhaps altered homeostatic regulation (Krishnan et al. 2011). Future MEA studies linking PLD1/PtdOH to network excitability patterns will be needed to address these intrinsic network and synaptic properties to maintain stable activity levels in response to chronic perturbations (Lezmy et al. 2020). Nevertheless, our MEA data uniquely demonstrates that the scope of PLD1 action extends beyond modulating the strength of individual synaptic connections (as measured by LTP magnitude in single-pathway recordings). Instead, it shapes the spatial organization and dynamics of activity across the network, coordinating activity among neuronal populations. This function is crucial for complex information processing, learning, and memory, which rely on precisely orchestrated network activity patterns. (Andrade-Talavera & Rodríguez-Moreno 2021). The finding that both OXP and ATT altered spatial patterns, even when ATT did not significantly change LTP magnitude, highlights that PLD1 modulation impacts how information flows and is represented across the hippocampal circuit. The increased prevalence of non-responsive sites and the induction of polarity flips with OXP, could signify a disruption of homeostatic plasticity crucial for maintaining network stability and preventing runaway excitation or quiescence (Lezmy et al. 2020), including altered oscillations and synchrony, that are increasingly recognized as a feature of AD contributing to cognitive impairment (Andrade-Talavera & Rodríguez-Moreno 2021). Lastly, even though PLD1 ATT did not significantly alter the magnitude of LTP in aged WT mice in single-electrode recordings, the MEA data revealed that it still exerted measurable effects by altering spatial response patterns and reducing paired-pulse facilitation (PPF). This demonstrates that reducing baseline PLD1 levels, even in the context of healthy aging, has subtle but detectable consequences for network function and presynaptic properties, thus reinforcing the notion that physiological levels of PLD1 activity (in a cell-type specific manner) are involved in the fine-tuning of network dynamics. Therefore, while reducing pathologically elevated PLD1 is a therapeutic goal, future studies evaluating the cell and brain region specific attenuation/overexpression and the relation to basal PLD1 activity will be instrumental in understanding the underlying mechanism important for synaptic resilience.

## EXPERIMENTAL METHODS

### Adeno-Associated Virus Production

Recombinant Adeno-Associated viral Vectors serotype 2 (AAV2) were constructed to modulate PLD1 expression in mouse models. The overexpression construct (PLD1 OXP) was created by inserting the full-length human PLD1 cDNA into the AAV vector plasmid, driven by a strong, constitutive promoter (specify the promoter, e.g., CMV, CAG). An enhanced green fluorescent protein (eGFP) cDNA was also included in this construct, enabling visualization of transduced cells. To attenuate mouse PLD1 expression, a short hairpin RNA (shRNA) sequence targeting mouse PLD1 (PLD1 ATT) was designed and cloned into the AAV vector. A control scrambled shRNA sequence (SCR), which does not target any known mouse mRNA, was used as a negative control, along with eGFP for visualization. All the constructs, except PLD1 OXP, included the eGFP gene. Recombinant AAV2 vectors (SCR/ATT and eGFP/OXP) were produced and titered by the University of North Carolina (UNC) Vector Core, ensuring high-quality and consistent viral preparations. The final viral titers were determined by qPCR and confirmed to be approximately 10_12_ pfu/mL. The Adeno-Associated Viral Vector Serotype 2 (AAV2) used in the current study is under the control of the CMV promoter. We and others have demonstrated higher specificity of AAV2 to neurons over astrocytes (Hammond et al. 2017; Haery et al. 2019).

### Animal Husbandry

This study was conducted in accordance with the National Research Council’s “Guide for the Care and Use of Laboratory Animals (8th Edition)” in the animal care facility at The University of Texas Medical Branch at Galveston (UTMB) which is accredited by the Association for Assessment and Accreditation of Laboratory Animal Care (AALAS), International. All procedures were approved by the Institutional Animal Care and Use Committee (IACUC) and were performed according to the National Institutes of Health (NIH) Guidelines on the use of laboratory animals. Male and female Wildtype (WT) mice, including C57BL/6J (RRID:IMSR_JAX:000664) and 129X1/SvJ (RRID:IMSR_JAX:000691) strains, and 3xTg-AD RRID:MMRRC_034830-JAX) transgenic mice were used in this study and maintained through a breeding program at UTMB. Mice aged 20 and 24 months were housed in social groups (3-5 mice per cage) under standard laboratory conditions: 23.3 ± 1.1°C temperature, 50% ± 20% humidity, and a 12-hour light/dark cycle. All mice received standard rodent chow and water ad libitum. Animal care and experimental procedures were performed in accordance with the guidelines set forth by the University of Texas Medical Branch (UTMB) Institutional Animal Care and Use Committee (IACUC) and adhered to national animal protection laws.

### Adeno-Associated Viral Vector Injections

Recombinant Adeno-Associated Viral vector Serotype 2 (rAAV2) was administered unilaterally into the cerebral lateral ventricle of mouse brains using stereotaxic surgery. Mice were anesthetized with isoflurane (1-3% in oxygen) and placed in a stereotaxic frame. A small burr hole was drilled into the skull at the following coordinates relative to bregma: anteroposterior (AP) −2.7 mm, lateral (L) +2.7 mm, and dorsoventral (DV) −3.0 mm. AAV vectors (SCR shRNA, PLD1 shRNA, PLD1 OXP, and eGFP) were injected using a Hamilton syringe at a volume of 3 µL and a titer of approximately 10^12 vg/mL. The injection rate was controlled to 0.5 µL/min to minimize tissue damage. Following injection, the burr hole was sealed with bone wax, and the scalp was sutured. Mice were monitored post-operatively until they recovered from anesthesia.

### Maxi-prep and Agarose Gel Electrophoresis

Plasmid DNA was purified using the Qiagen EndoFree Plasmid Maxi Kit (Cat. no. 12362) according to the manufacturer’s protocol. Briefly, bacterial colonies from glycerol stocks of pAAV.PLD1.MCS (OXP) and pAAV.PLD1.shRNA (ATT) were grown in 3 mL of Miller’s LB broth supplemented with ampicillin (100 μg/mL) at 37°C for 6-8 hours. This culture was then transferred to 250 mL of LB broth with ampicillin and incubated overnight at 37°C with shaking (225 rpm). Bacterial cells were harvested by centrifugation (5,000 rpm, 15 min, 4°C), and the resulting pellet was resuspended in TE buffer (pH 8.0).

Restriction enzyme digestion and agarose gel electrophoresis were performed to confirm the integrity and size of the plasmid constructs. Double digestion of the plasmids (OXP and ATT) was carried out using PvuII-HF (NEB, Cat. no. R3151) and PstI-HF (NEB, Cat. no. R3140) restriction enzymes, following the manufacturer’s instructions. The digested fragments were then separated by electrophoresis on a 1% TAE agarose gel, along with an appropriate DNA ladder (Ready-to-Load High Molecular Weight DNA Ladder, with eleven bands ranging from 0.5 kb to 12 kb).

### Animal Behavior Assessment

#### Y-maze test (1 day)

Spatial working memory was assessed using a symmetrical Y-shaped maze (San Diego Instruments). Mice were placed in one arm and allowed to explore for 8 minutes. Arm entries and sequences were recorded. Spontaneous alternation was defined as consecutive entries into all three arms. Animals with fewer than five entries were excluded. Total entries and latency to exit the starting arm were also recorded. Successful alternations included three consecutive visits to a different arm, i.e., the successive arm was not visited immediately prior to the current arm. Each animal was placed in a symmetrical Y-shaped maze (arm dimensions: length - 40cm; width - 8cm; height - 12cm (San Diego Instruments) that was beige colored with non-reflective surfaces. Each mouse was placed in any one arm while facing the center (designated as arm A) and allowed to explore the maze for 8 minutes. The arm to the right of this arm was B and the arm to the left was C. Arm A was varied for each animal to prevent bias of arm placement. Latency to leave the first arm and total number and sequence of entries into each arm were scored for each mouse. An arm entry was accounted when the mouse had all four paws inside the arm. Animals making fewer than five entries were not counted in the final analyses – they were excluded as non-participators. A spontaneous alternation was defined as successive entries into each of the three arms on overlapping triplet sets (e.g., ABC, BCA, CAB, etc.). Percentage of spontaneous alternation performance (%SAP) was calculated as the ratio of actual alternations (total alternations) to possible alternations (total arm entries − 2) × 100. In addition, total entries were scored as an index of locomotor activity and the latency to exit the starting arm as emotionality-related behavior. After the 8-minute test, the animal was returned to its home cage and allowed to rest for 24 hours before the Novel Object recognition paradigm.

#### Open Field Test (OFT) and Novel Object Recognition (NOR)

Mice were acclimated to the open field arena over three 15-minute sessions, 24 hours apart. Locomotor activity was quantified using TopScan software (Clever Sys. Inc.). For NOR, mice were exposed to two identical objects for 10 minutes (training). After a retention interval (2-24 hours), mice were exposed to one familiar and one novel object for 10 minutes. Object exploration time was recorded using ObjectScan software (Clever Sys. Inc.). Each mouse was habituated to test for normal locomotion and acclimation to the test environment. After placement in the open field box for two 10-minute test sessions that were 24 hours apart, the TopScan (Clever Sys. Inc.) video tracking software quantified various locomotor parameters: total distance traveled, time spent moving >50mm/sec, number of rears, number of entries into and time spent in the center 1/9th of the locomotor arena. The data collected during the habituation phase was used to assess the OFT outcomes. The amount of time spent in different zones (edges vs central) was assessed every day. Twenty-four hours after the last habituation session, mice were subjected to training in a 10-minute session of exposure to two identical, non-toxic objects (metal or hard plastic items) in the open field box. The time spent exploring each object was recorded using ObjectScan (Clever Sys. Inc.); an area 2 cm² surrounding the object was defined such that nose entries within 2 cm of the object were recorded as time exploring the object. After the training session, the animal was returned to its home cage. After a variable retention interval of 2 minutes to 24 hours, the animal was returned to the arena in which two objects, one identical to the familiar object but previously unused (to prevent olfactory cues and prevent the necessity to wash objects during experimentation) and one novel object. The animal was allowed to explore for 10 minutes, during which the amount of time exploring each object was recorded. Objects were randomized and counterbalanced across animals. The animals were returned to their home cages with food/water ad-libitum for a minimum of 24 hours.

#### Elevated Plus Maze (EPM)

The EPM evaluates exploratory behavior in a novel and anxiety-inducing environment. The apparatus consisted of two closed arms and two open arms, each measuring 12 × 50 cm, elevated 75 cm above the floor. Photobeams were positioned at the entrance of each arm (Med Associates Inc, VT, USA) to monitor the time spent on the open arms. The mice’s movements were tracked for 5 minutes using Med-PC software, which recorded photobeam interruptions. Mice were placed in the maze for 5 minutes, and time spent in open and closed arms was recorded using video tracking software.

#### Forced Swim Test (FST)

The FST was used to assess depression-like behavior, but only for WT mice and only at 20-months given the level of stress associated. Mice were placed into a 2-liter beaker filled with approximately 1.2 liters of room temperature water (24 ± 0.5°C) for 15 minutes on the first day (Session 1) and for 5 minutes on the second day (Session 2). After each session, the mice were dried and returned to their home cages. Swimming activity was video recorded, and the latency to the first period of immobility (defined as 1 second) and total time immobile were determined for Session 2 by an investigator blinded to the conditions.

#### Fear Conditioning (FC)

Associative learning and memory were assessed using fear conditioning. The standard protocol consisted of a training phase where the mouse was placed in a particular environment (a training chamber with specific lighting, geometry, and odor) and allowed to explore for three minutes. An auditory stimulus (80dB white noise) was then presented for 30 seconds. This was the auditory conditioned stimulus (CS). Additionally, there was the context CS comprised of visuospatial, olfactory, and auditory cues. One foot shock (0.5 mA, two seconds duration; this was the unconditioned stimulus, US) was delivered during the last two seconds of the auditory CS. A second presentation of the auditory CS and the US was delivered at the five-minute mark, and the animal was then left in the cage for another 2.5 minutes. Twenty-four hours later, the animal was returned to the same training chamber, and the context test for fear learning was performed. The amount of freezing the rodent exhibited during five minutes in the training chamber was measured. One hour later, the cued test was performed in a completely novel context. The animal was placed in the testing chamber, and freezing was measured for three minutes. Then the auditory CS was presented again, and freezing was quantified over the next three minutes. The animals were returned to their home cages with food and water ad libitum for 24 hours.

### Field Electrophysiological Recordings

#### Conventional Electrophysiology

Our standard protocol was used as previously described (Bourne et al. 2019; Natarajan et al. 2023). Briefly, mice were deeply anesthetized with isoflurane and transcardially perfused with ∼30 mL of room temperature carbogenated (95% O_2_ and 5% CO_2_ gas mixture) NMDG-artificial cerebrospinal fluid (aCSF) (in mM - 93 N-Methyl-D-Gluconate, 2.5 KCl, 1.2 NaH_2_PO_4_, 30 NaHCO_3_, 20 C_8_H_18_N_2_O_4_S, 25 C_6_H_12_O_6_, 5 C_6_H_7_O_6_Na, 2 CH_4_N_2_S, 3 C_3_H_3_NaO_3_, 10 MgSO_4_,7H_2_O, 0.5 CaCl_2_,2H_2_O, 12 C_5_H_9_NO_3_S, pH 7.4), and sliced using Compresstome VF-300 (Precisionary Instruments, Greenville, NC) in carbogenated NMDG-aCSF to obtain 350 μm transverse brain sections. Slices were allowed to recover for 10 min in carbogenated NMDG-aCSF at 33°C. Slices were then maintained at room temperature in a modified carbogenated HEPES holding aCSF solution (in mM - 92 NaCl, 2.5 KCl, 1.2 NaH_2_PO_4_, 30 NaHCO_3_, 20 C_8_H_18_N_2_O_4_S, 25 C_6_H_12_O_6_, 5 C_6_H_7_O_6_Na, 2 CH_4_N_2_S, 3 C_3_H_3_NaO_3_, 2 MgSO_4_,7H_2_0, 2 CaCl_2_,2H_2_0, 12 C_5_H_9_NO_3_S, pH 7.4). Slices were recorded in carbogenated standard recording NaCSF (in mM - 124 NaCl, 2.5 KCl, 1.2 NaH_2_PO_4_, 24 NaHCO_3_, 5 C_8_H_18_N_2_O_4_S, 13 C_6_H_12_O_6_, 2 MgSO_4_,7H_2_0, 2 CaCl_2_,2H_2_0, pH 7.4). Evoked field excitatory post-synaptic potential (fEPSP) recordings were performed by stimulating the Schaffer collateral pathway (located in stratum radiatum) using a stimulating electrode of ∼22 kΩ resistance placed in the CA3 region and glass recording electrodes in the CA1 region. Current stimulation was delivered through a digital stimulus isolation amplifier (A.M.P.I, ISRAEL) and set to elicit a fEPSP approximately 30% of maximum for synaptic potentiation experiments using platinum-iridium tipped concentric bipolar stimulating electrodes (FHC Inc., Bowdoin, ME). The use of platinum iridium wire and diphasic pulses can help minimize electrode polarization (Mathis et al. 2011). Using a horizontal P-97 Flaming/Brown Micropipette puller (Sutter Instruments, Novato, CA), borosilicate glass capillaries were used to pull recording electrodes and filled with NaCSF to get a resistance of 1–2 MΩ. Field potentials were recorded in CA1 stratum radiatum using a Ag/AgCl bridge with CV7B headstage (Molecular Devices, Sunnyvale, CA) located ∼1–2 mm from the stimulating electrode. LTP was induced using a high frequency stimulation protocol (3 x 100 Hz, 20 seconds) as previously described (Bourne et al. 2019; Natarajan et al. 2023). To assess basal synaptic strength, 250 μs stimulus pulses were given at 10 intensity levels (range, 100–1000 μA) at a rate of 0.1 Hz. Three field potentials at each level were averaged, and measurements of fiber volley (FV) amplitude (in millivolts) and fEPSP slope (millivolts per millisecond) were performed using Clampfit 10.7 software. Synaptic strength curves were constructed by plotting fEPSP slope values against FV amplitudes for each stimulus level. Baseline recordings were obtained for 10 minutes by delivering single pulse stimulations at 20 second intervals. All data are represented as a percentage change from the initial average baseline fEPSP slope obtained for the 10 min prior to HFS. Maximum of two slices were recorded per animal and averaged to give the response per animal.

#### Multi-electrode Array (MEA)

Hippocampal slices generated using procedures specified above were positioned on a Multi-Electrode Array (MEA) to study neural network activity. The slices were placed on the MEA, ensuring good contact with the electrodes, and a perfusion system was established to continuously exchange artificial cerebrospinal fluid (ACSF, Base A + Base B, #LRE-S-LSG-1000-1, EcoCyte Bioscience) and provide oxygenation and temperature control. Electrical stimuli were applied using one or more electrodes to evoke activity in the slices, with stimulation parameters adjusted to study different aspects of neural network function. The MEA recorded the electrical activity of the neurons, which was analyzed to identify synaptic potentials and population spikes. MEA recordings were performed using a MED64 system (Alpha MED Sci) to assess synaptic function in the CA1-dentate gyrus region. Input-output curves, paired-pulse facilitation, and long-term potentiation (LTP) were measured following electrical stimulation. Data were digitized and analyzed using Mobius software.

### Cell Culture

Rat pheochromocytoma (PC12) cells were maintained in RPMI 1640 medium supplemented with 5% fetal bovine serum and 5% horse serum. For NGF-induced differentiation, cells were treated with NGF (100 ng/mL) under reduced serum conditions for 7 days. Brightfield and fluorescence images were captured using a Leica microscope.

### Statistical Analysis

All data were reported as mean ± SEM. Statistical significance was calculated using GraphPad Prism 9.2 (San Diego, CA). All statistical tests were two-tailed, with the threshold for statistical significance set at 0.05. To account for non-normal distribution of data, either non-parametric t-tests (Mann-Whitney U or Wilcoxon rank sum) or one-way ANOVA (Kruskal-Wallis test) followed by Geisser-Greenhouse correction for mixed effects analysis, Holm-Sidak multiple comparisons test with individual variances computed for each comparison, or uncorrected Fisher’s LSD with individual variances computed were applied as appropriate to account for variability of differences. Double blinding was performed by having one scientist perform the dilutions and provide them to the experimenter conducting the injections and subsequent experiments with a code denoting the different treatments. Once the analysis was completed, the code was broken. Statistical analysis was performed using GraphPad Prism 10, with two-tailed t-tests (Mann-Whitney U) or one-way ANOVA (Kruskal-Wallis) followed by Dunn’s multiple comparisons used to determine statistical significance. A p-value of 0.05 was considered significant.

## ACKNOWLEDGEMENTS

This work was supported by grants from the Jeanne B. Kempner Scholarship (2022-2023, C.N.), AARG-17-533363 (B.K.), NIA R01 – AG063945 (B.K.), The Don and Nancy Mafrige Professor in Neurodegenerative Disease Endowment (B.K.), the Mitchell Center for Neurodegenerative Diseases, and NIA R21 - AG059223 (B.K.). The authors gratefully acknowledge Dr. John Allen and his graduate student, Ryan Murphy, for their assistance in characterizing the plasmids, and the University of North Carolina (UNC) Vector Core for their expert packaging of the plasmids into AAV2 vectors. We thank Dr. Giulio Taglialatela and his laboratory members for their insightful feedback and constructive criticism of the data.

## DATA AVAILABILITY STATEMENT

The data that support the findings are available from the corresponding author upon reasonable request.

## AUTHOR CONTRIBUTIONS

SGS: Data curation, Investigation, Methodology, Writing-original draft; SMB, SM, MEV, PS, SKJ, CN, KR, KS, EV, PCP, JLL, SKG, JC, KG : Data curation, Investigation Methodology; AL, TAG: Data curation, Software, review and editing; BK: Conceptualization, Data curation, Funding acquisition, Investigation, Methodology, Project administration, Resources, Software, Supervision, Writing – review and editing.

---

**Schematic Figure:**
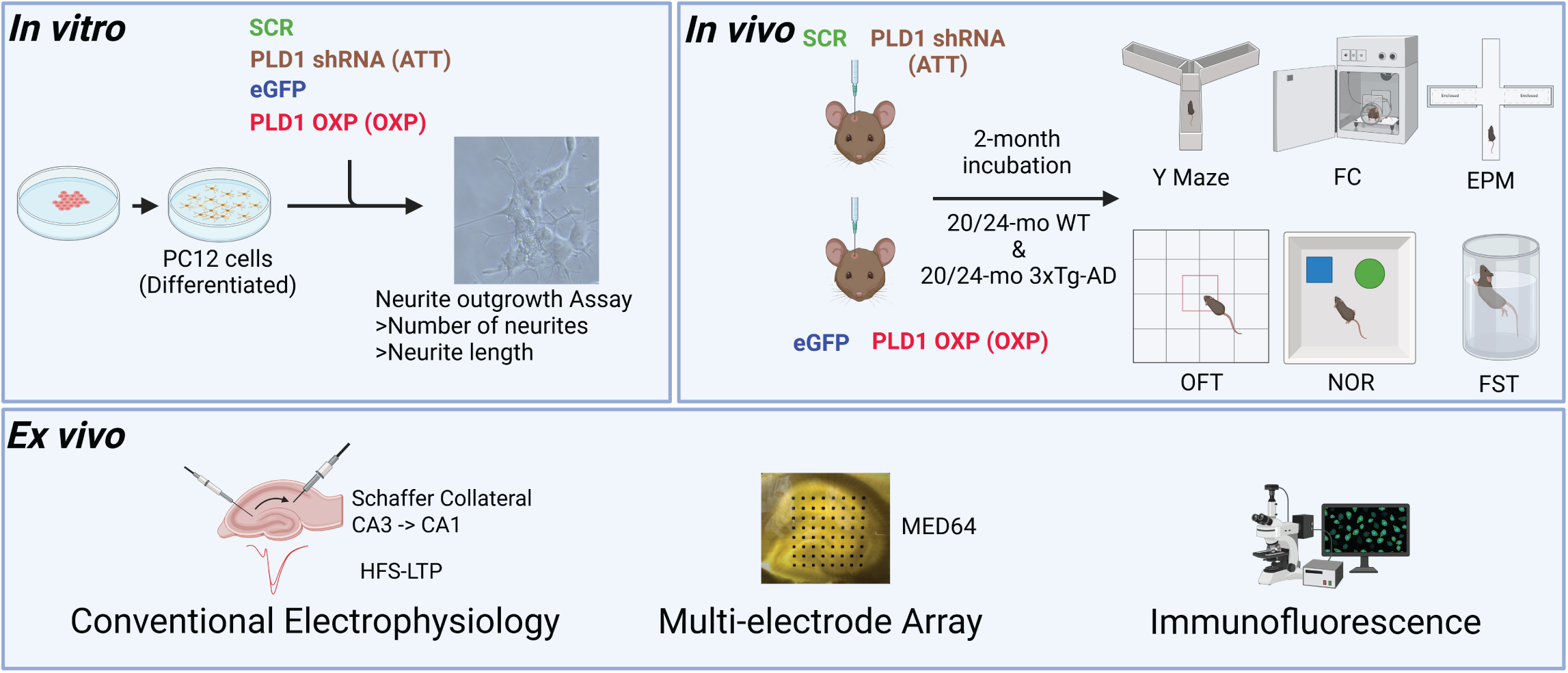

**Supplementary Figure S1:**
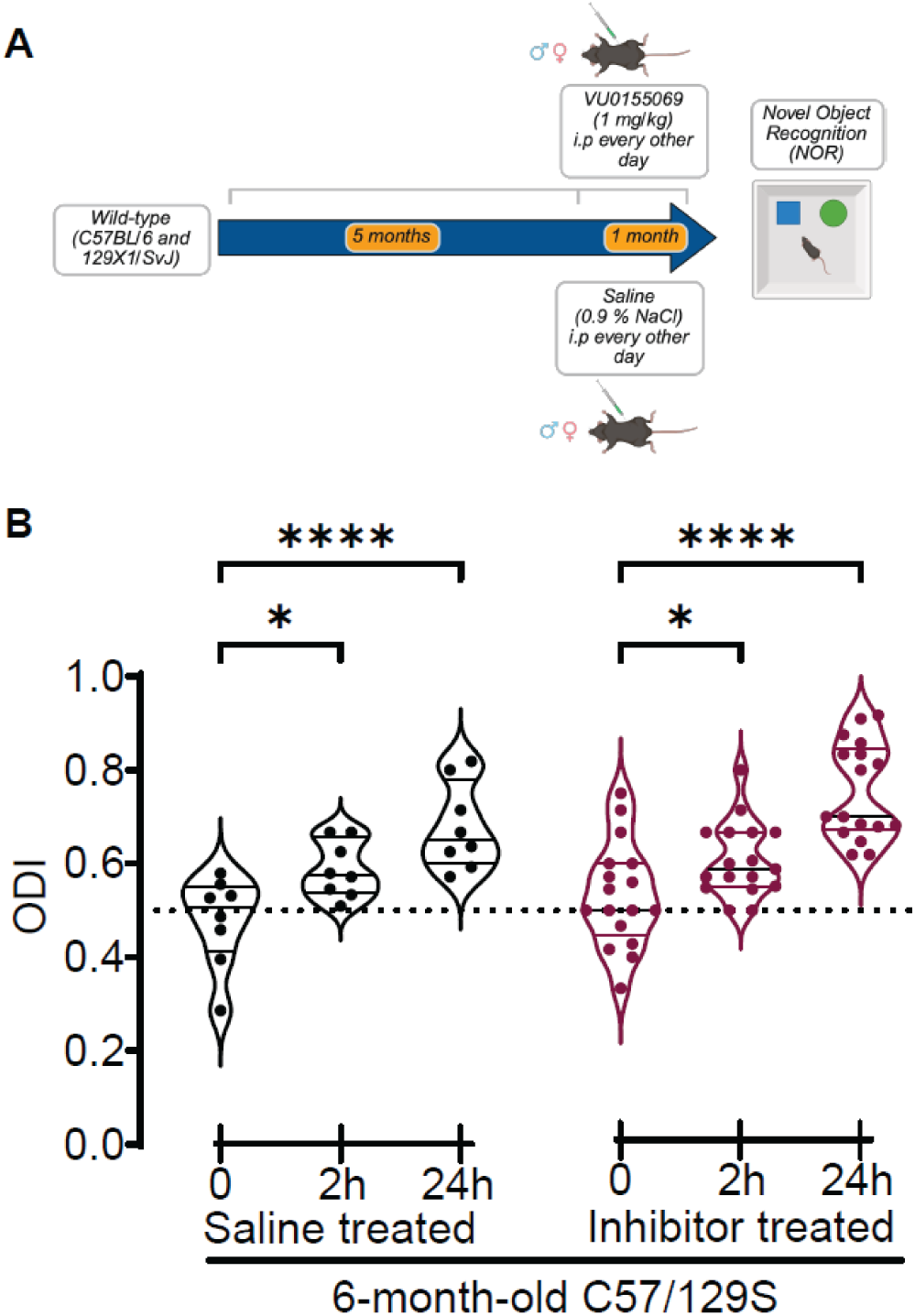

**Supplementary Figure S2:**
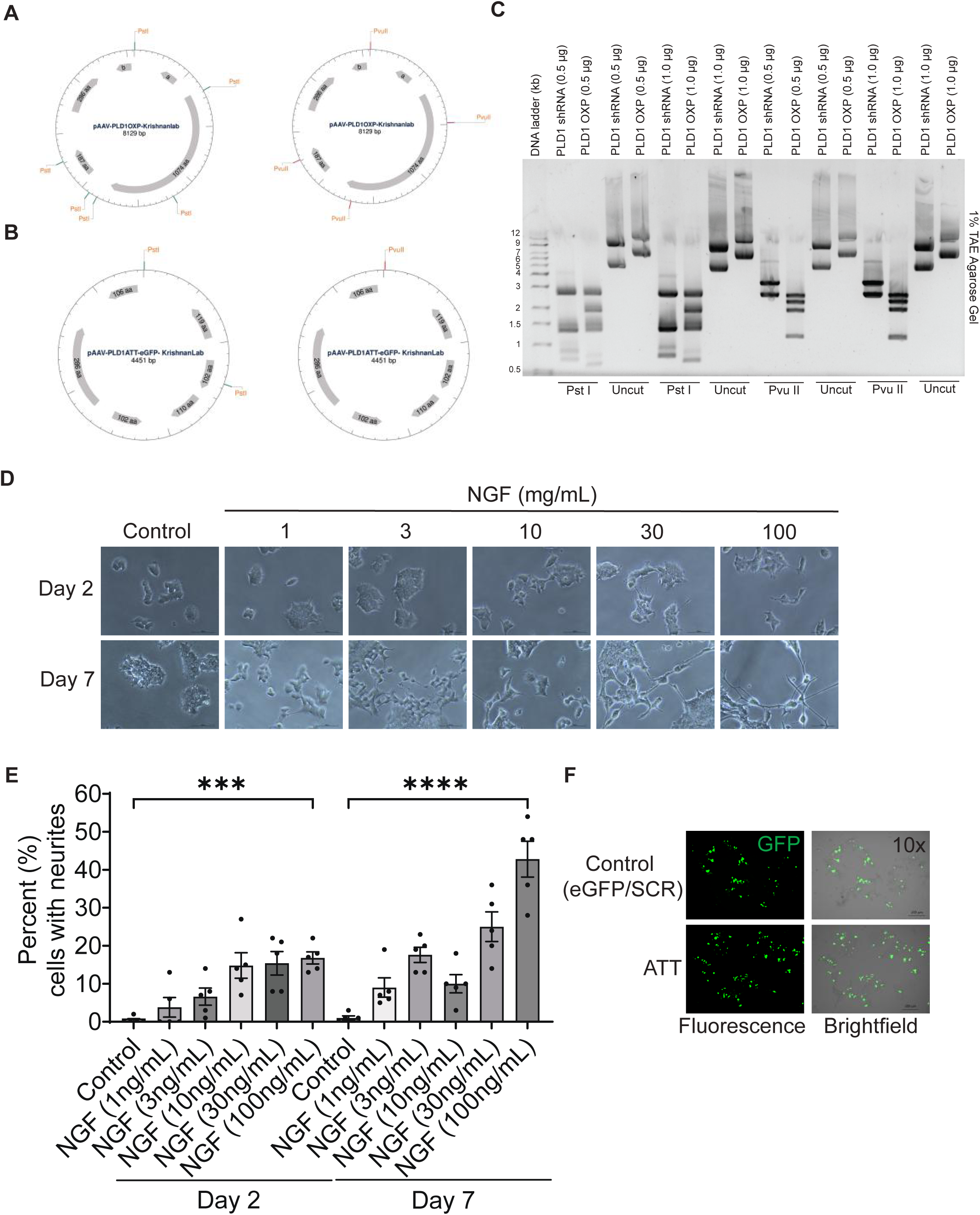

**Supplementary Figure S3:**
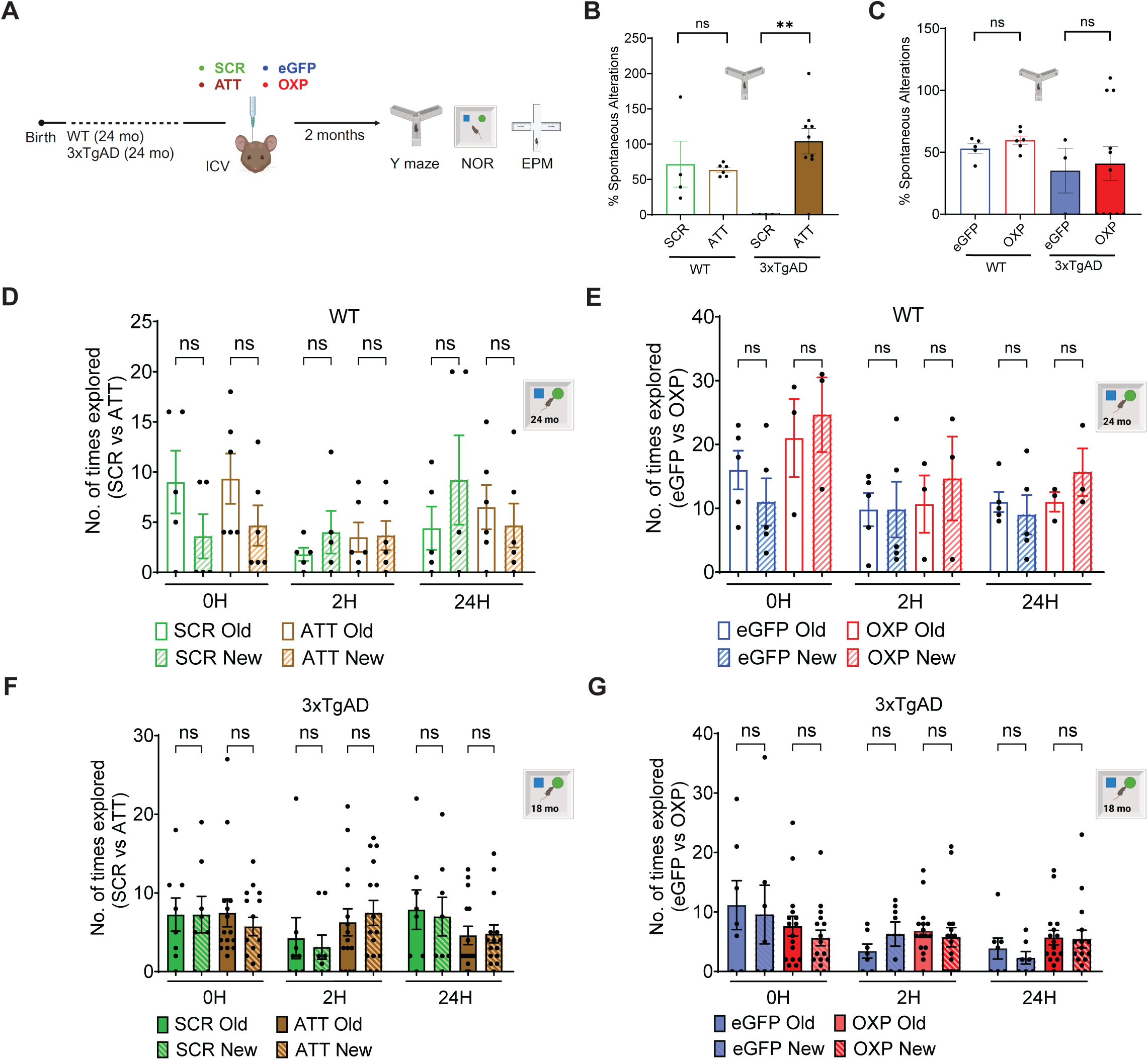

**Supplementary Figure S4:**
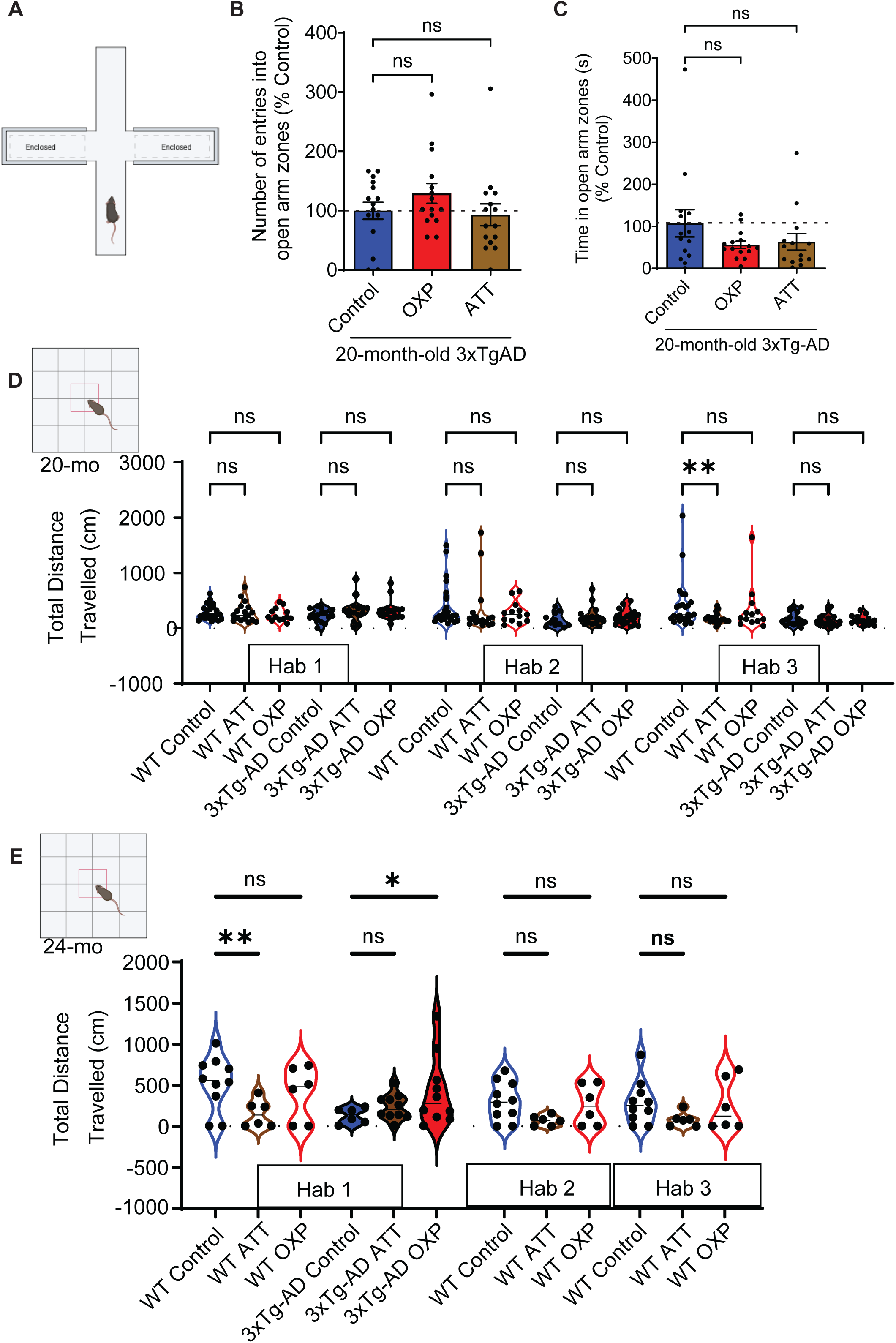

**Supplementary Figure S5:**
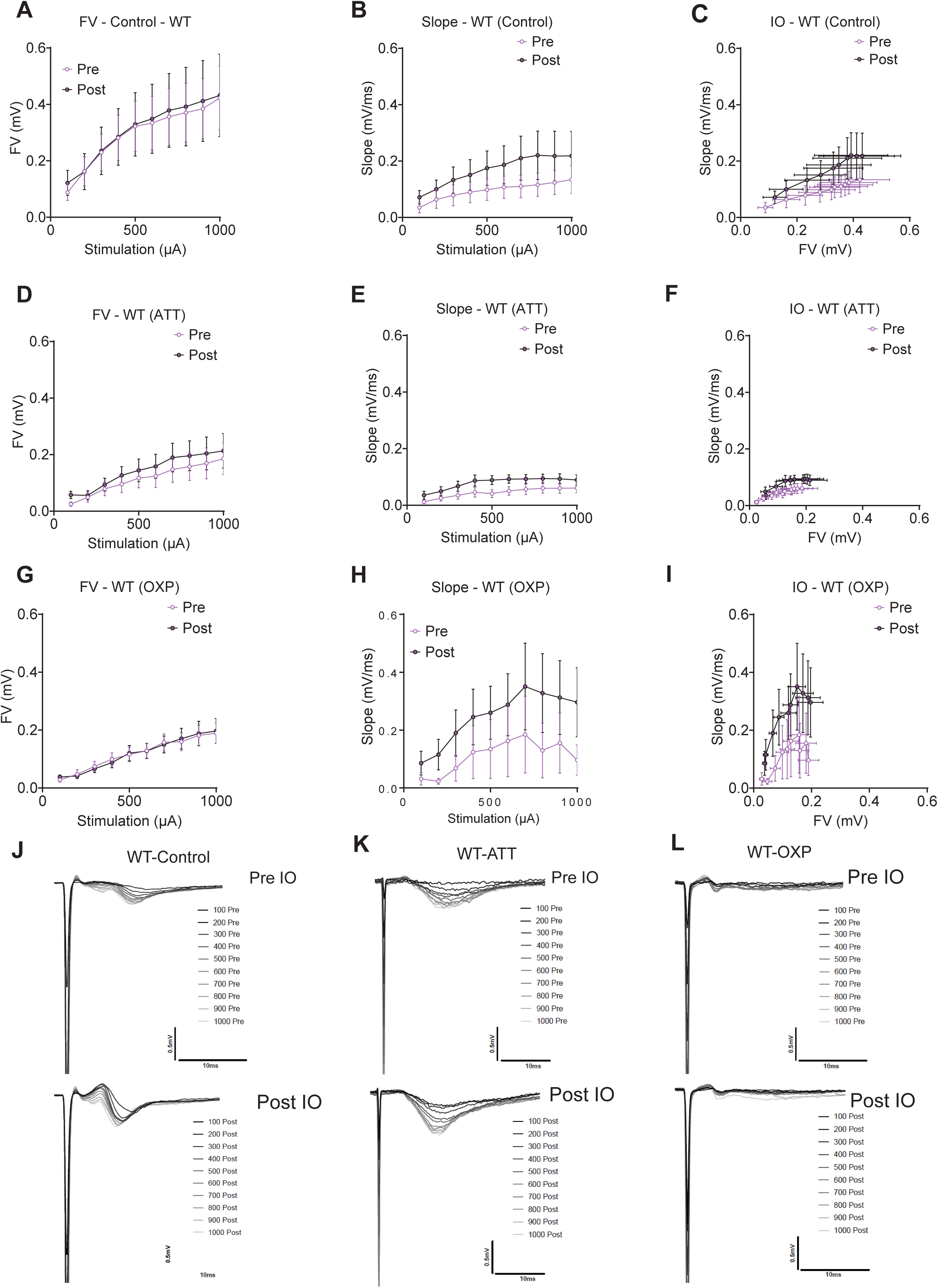

**Supplementary Figure S6:**
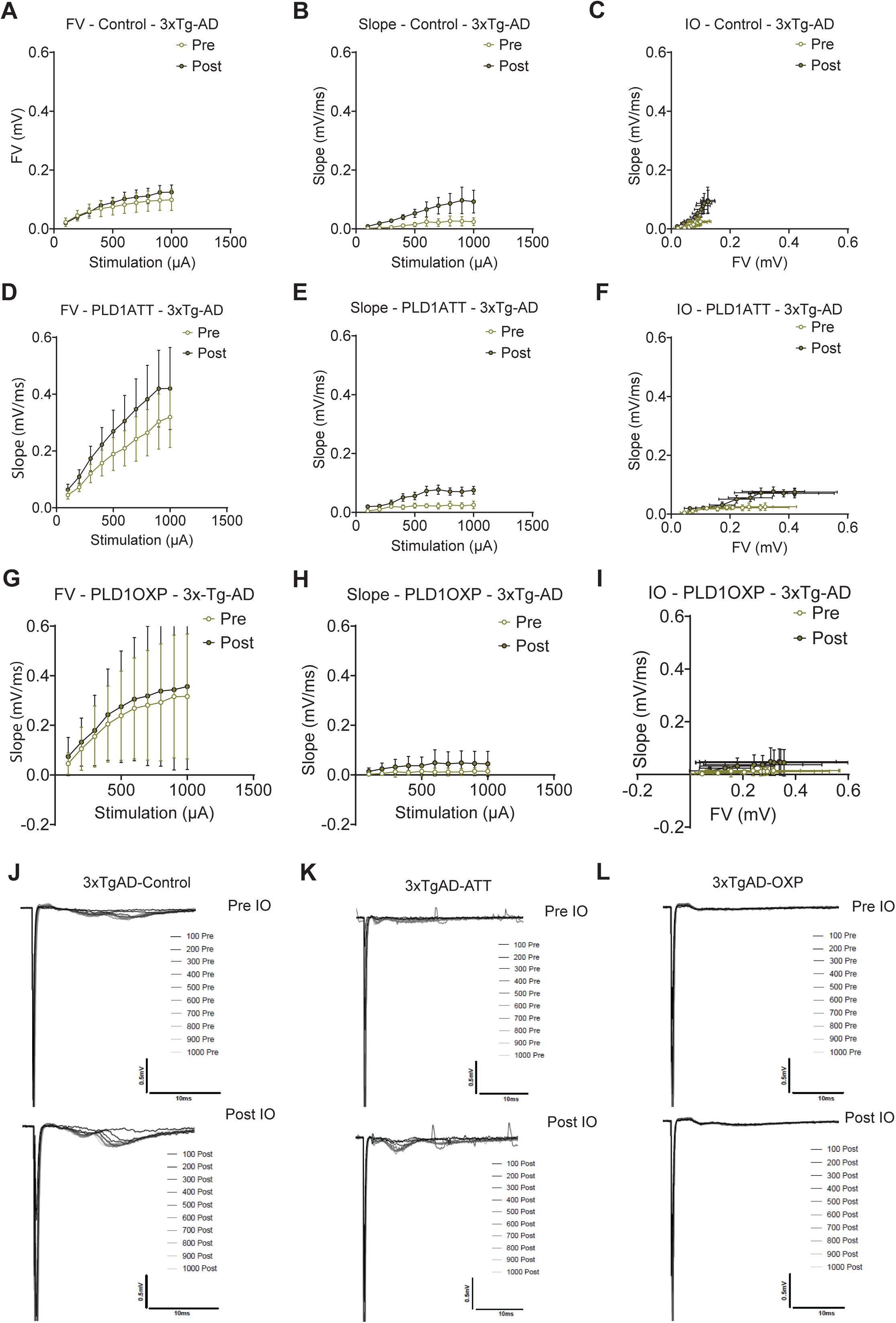

**Supplementary Figure S7:**
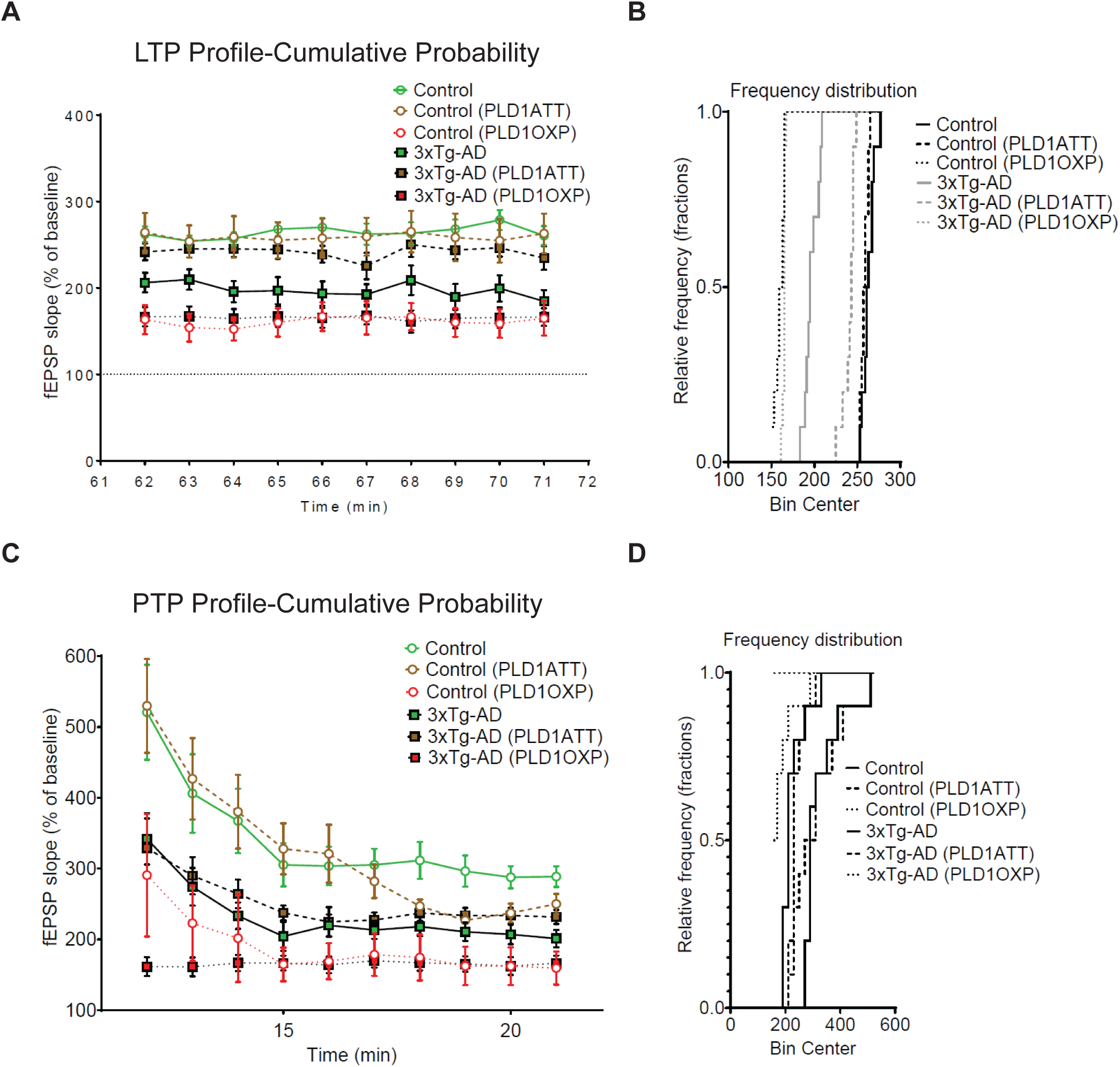

**Supplementary Figure S8:**
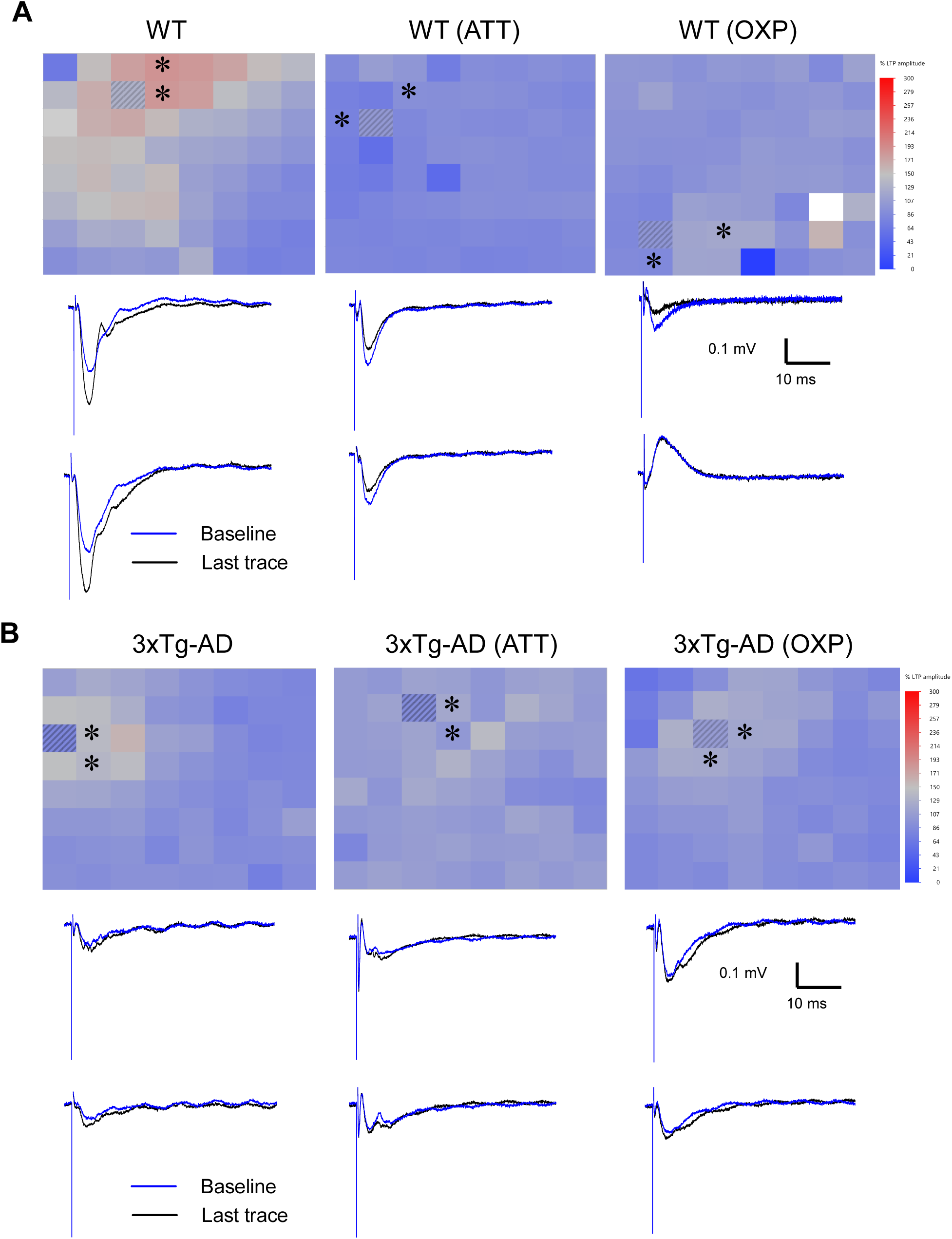

**Supplementary Figure S9:**
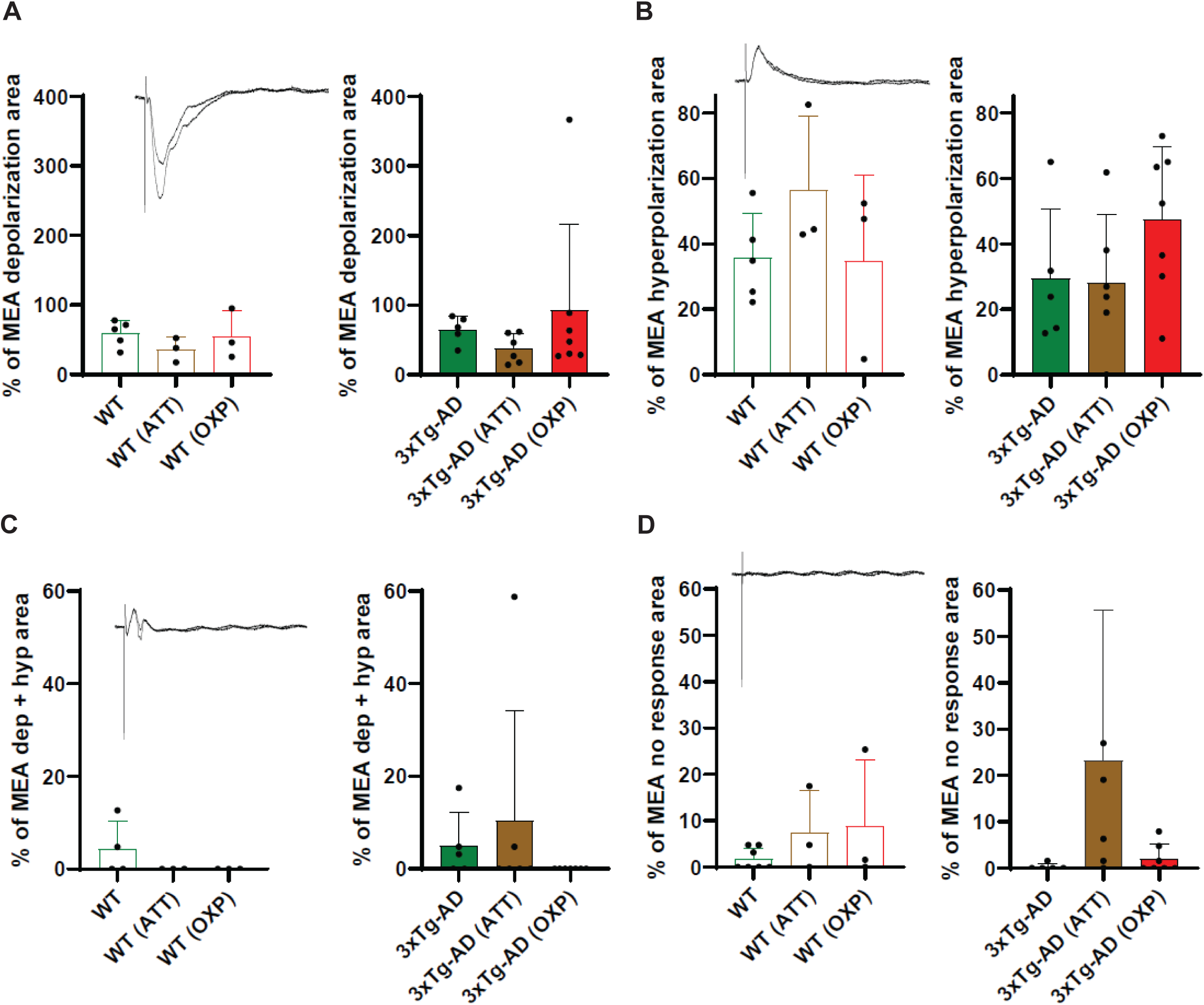

**Supplementary Figure S10:**
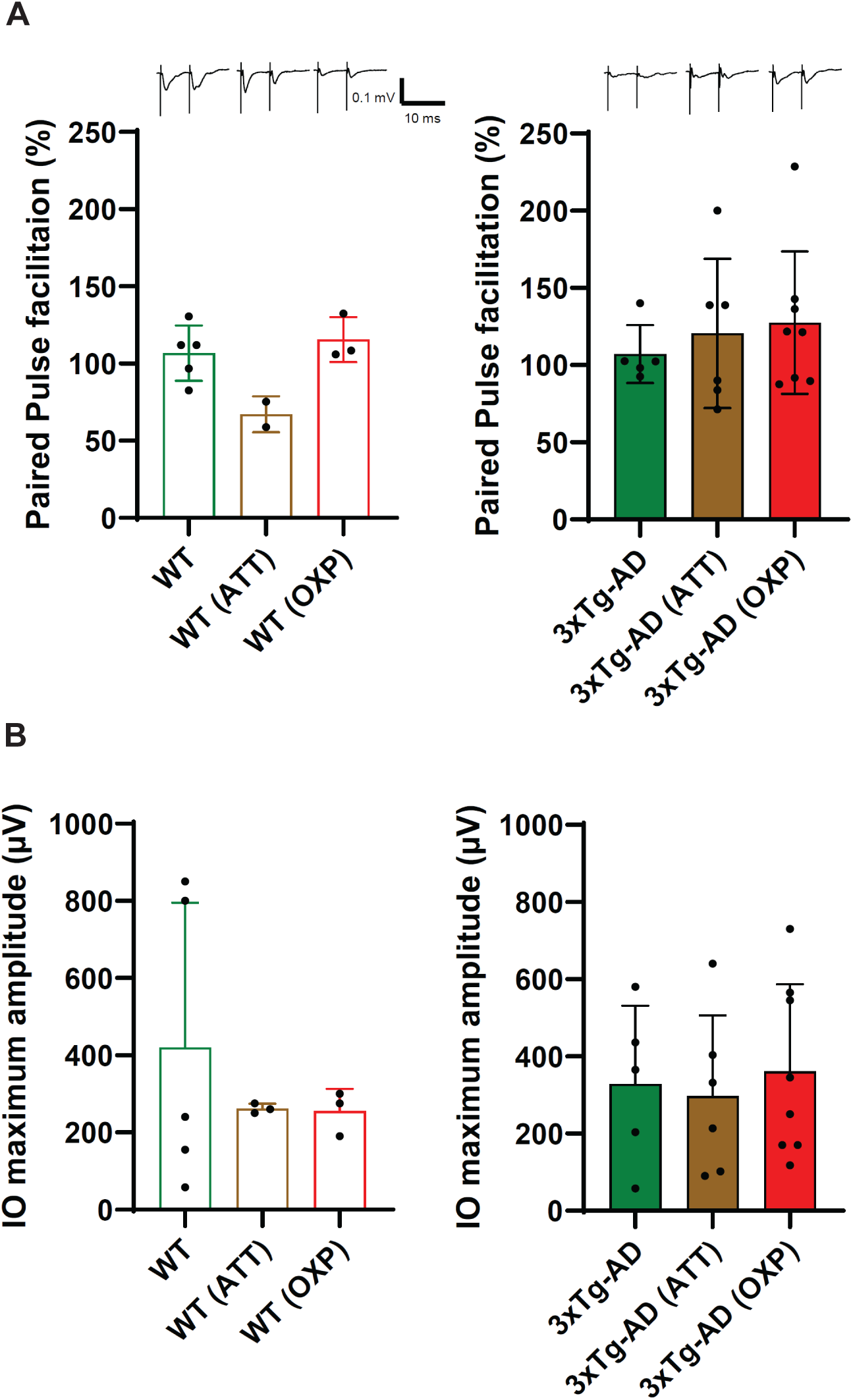

**Supplementary Figure S11:**
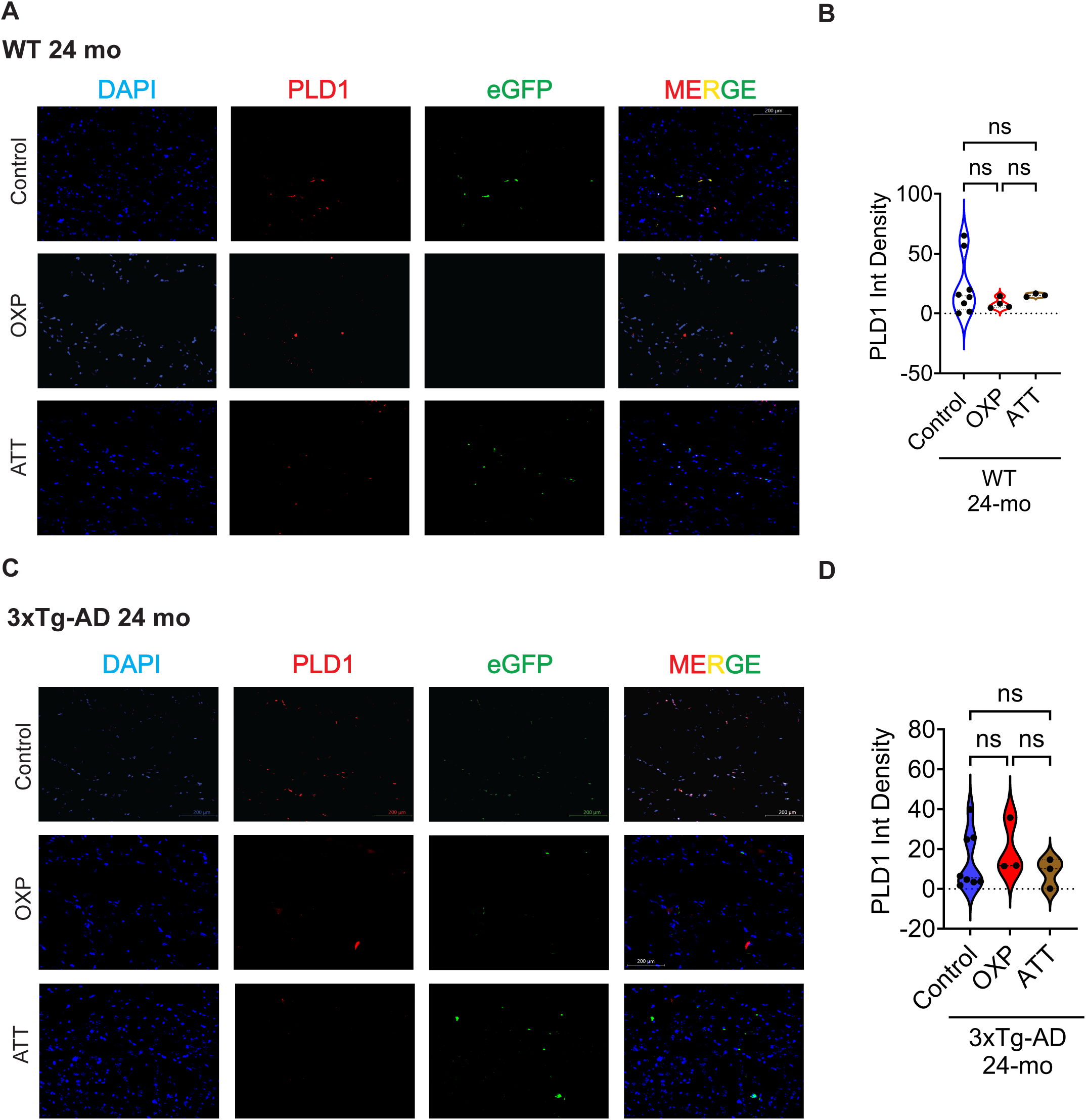

**Supplementary Figure S12:**
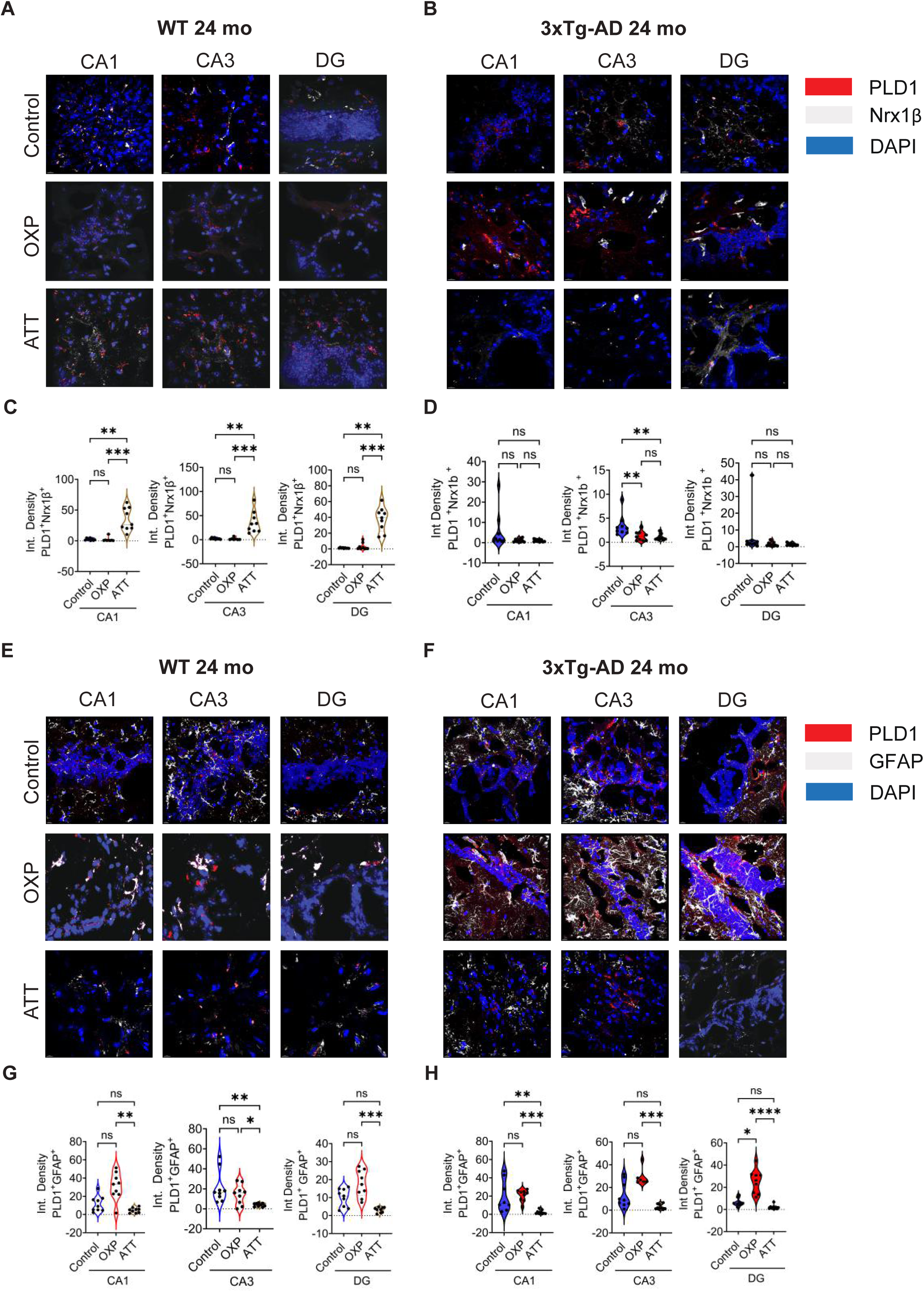

